# Next-Generation Breast Organoids Capture Human Organogenesis with High-Resolution Live Imaging

**DOI:** 10.1101/2023.10.02.560364

**Authors:** Gat Rauner, Nicole C. Traugh, Colin J. Trepicchio, Meadow E. Parrish, Kenan Mushayandebvu, Charlotte Kuperwasser

## Abstract

Organoids have emerged as a powerful tool for modeling tissue growth and diseases. In this study, we introduce a groundbreaking organotypic culture technique that replicates the morphology, scale, and heterogeneity of human breast tissue, and includes a mesenchymal-like stromal component. A standout feature of this approach is the use of long-term live imaging at high temporal resolution to directly observe stem cell dynamics during organogenesis, from single cells to mature organ tissue. The system is adaptable for high throughput applications and allows for genetic manipulation of the cells. Real-time imaging of ex-vivo tissue formation reveals a non-canonical process of ductal-lobular morphogenesis and branching, and de-novo generation of a supportive stroma. Incorporating patient-derived single cells from multiple donors offers an enhanced representation of the spectrum of individual responses and the impacts of distinct exposures. While developed for breast tissue, the principles of this technology can serve as a model for the development of similar systems in other tissues, where organoids do not merely reproduce the tissue, but where their regeneration can also be observed and studied. In addition, this model provides a quantitative experimental system to study mechanisms of embryogenesis, development, and tissue organization where biomechanics plays an important role.

## Introduction

Organogenesis, including tissue morphogenesis, is essential in the development and maintenance of multicellular organisms. This process involves the coordination of cell behavior and organization, allowing for the formation of functional tissues and organs. Understanding how tissues develop and form is crucial for understanding how they work and how they can be repaired or regenerated in the case of disease or injury. In addition, defects in organogenesis can lead to developmental disorders and birth defects, and insights into the cellular and molecular mechanisms that are disrupted can potentially lead to new treatments or preventative measures. Finally, understanding how cells communicate and work together to form complex structures and systems has implications for a wide range of fields beyond developmental biology, including tissue engineering and regenerative medicine.

Organoids are miniaturized and simplified versions of organs produced in vitro in three dimensions that show realistic micro-anatomy. They are derived from stem cells and can replicate some of the organ’s structure and function. Human breast organoids have emerged as a promising tool for studying developmental processes and breast cancer ^1–3^. However, these organoids are mostly spheroid and rudimentary, and thus are unable to establish physiologically relevant complex patterning, hormonal sensitivity, scale, or relevant architectural complexity. Most traditional breast organoid models rely on an extracellular matrix scaffold composed entirely from Matrigel or Type I Collagen, which only provides a supportive microenvironment for cells to promote self-assembly of simple and rudimentary structures ^4–9^. Moreover, the focus of traditional organoid models has largely been on the prominent parenchymal cells, often the epithelium, overlooking the supportive stromal cells, which provide structural support and are crucial for organ functionality. These stromal cells not only provide structural support but also partake in essential interactions that influence tissue development and hormonal responsiveness. Thus, traditional breast organoid models have had limited value in studying tissue morphogenesis and disease, including cancer, as they lack many key features of the organ.

Development of improved biomimetic ECM scaffolds is an active area of research, with the goal of providing more accurate and reproducible models of in vivo tissue microenvironments. Recently, collagen-based hydrogels containing components that are part of the human breast ECM: fibronectin, laminin, and hyaluronic acid, have been reported ^10^. Here, we used this newly described biomimetic ECM scaffold to develop an organoid technology that not only recapitulates in vivo tissue, but also serves as an observational model of organ development. Notably, human organoids cultured in this ECM do not require “niche factor” media supplements such as Wnt3a, Noggin or R-spondin, or inhibitors of ROCK, TGF-β or MAPK signaling ^11–13^. This technology can therefore facilitate fundamental studies of development, as well as potential therapeutic interventions.

From a single adult human breast stem cell, a miniaturized human breast is generated, comprised of several terminal ductal-lobular units (TDLUs). To account for human patient variability, we generated organoids from multiple human donors, and analyzed their frequency and regenerative capacity. We used single-cell RNA-seq to demonstrate that the organoids recapitulate multiple cell populations found in the human breast tissue, including a stromal-like population. By adapting organoid culture to high-throughput, long-term confocal live-imaging microscopy, we gain an unprecedented temporal resolution view, in real time, of organogenesis, depicting the cell dynamics from single cells to complex tissue, over 18 days. Importantly, we demonstrate the capability of this model to support genetic modification of primary cells using lentiviral transduction, enabling the use of genetic reporters, screening, and other tools, in the context of human tissue development. Moreover, we combine live-imaging microscopy with a sophisticated tracking algorithm quantify cell velocity, directionality, and displacement, as well as organoid over time in a sequence of videos.

We find that organoid formation proceeds through distinct phases: During the first 4–7-days cells go through a quiescent phase of induction, followed by an active patterning phase in which detached motile cells trace the “blueprint” of the future branching pattern. The patterning phase is followed by a morphogenesis phase, in which parenchymal cells proliferate and populate the pre-determined branching blueprint, giving rise to the ductal-lobular architecture. Concomitantly, populations of stromal-like cells emerge from the growing organoid, proliferate and migrate throughout the hydrogels. Lastly, in the third week of culture, the established ductal-lobular organoid matures to its final TDLU morphology.

The branching process we observe and describe here is in striking contrast to the well characterized process of budding, where a terminal end-bud invades the matrix. Furthermore, we observe the de-novo creation of a mesenchymal-like stroma that surrounds the developing organoid parenchyma that can be seen pulling and pushing the surrounding ECM, contributing to the shaping and bending of the developing breast organoid tissue

We apply this enhanced organoid technology to evaluate organoid formation from cells obtained from pre-menopausal and post-menopausal women, to study the effects of hormonal changes in vivo on organogenesis and regeneration capacity ex-vivo. Together, these findings describe a robust method for generating, studying, and tracking patient-derived biomimetic breast tissue, that can be used for clinical and basic biology studies.

## Design

Recognizing the limitations of current breast organoid models, and the pressing need for physiologically relevant models that could mimic tissue morphogenesis more accurately, we sought to design a biomimetic model that would not only replicate its in vivo counterpart, but also serve as a window into bona-fide human organogenesis. The design considerations were:

### Composition of the ECM scaffold

Rather than solely relying on Matrigel or Type I Collagen, it was important to integrate additional components native to the human breast ECM. Recent literature highlighted the role of fibronectin, laminin, and hyaluronic acid ^10^. These components were considered essential in the design to ensure the scaffold could mimic the native breast tissue environment more closely.

### Faithful tissue representation

For the design to offer genuine insights and applications, it was essential to ensure that the multitude of cell types present in vivo is accurately recapitulated in the model, leading to more authentic cellular behaviors, interactions, and responses within the organoid model.

### Observation at single-cell level

Starting with single cells embedded within the ECM enables detailed tracking of the initial stages of cell differentiation and proliferation, offering insights into the nuances of early organogenesis.

### High-resolution, real-time tracking

An integral part of this design was to enable real-time tracking of organoid development. The design incorporated high-throughput, long-term confocal live-imaging microscopy as well as lentiviral transduction of fluorescent reporters and algorithms to quantify and track cell movement and organoid formation to provide an uninterrupted observational window into the entire organ development. This enables further insights into the organogenesis processes.

### Accurate representation of in vivo variation

The design aimed to capture the inherent variability among humans by utilizing patient-derived cells from diverse donors, reflecting differences due to factors like age and hormonal exposure.

The resulting design centered around donor-derived human breast stem cells embedded within a conducive ECM. This setup, when combined with an advanced imaging system, permitted unparalleled observation of the entire spectrum of organogenesis, from the earliest cellular events to tissue maturation.

## Results

### Regeneration, architecture, and patterning of patient derived organoids from single cells

Organoids were generated from primary single cells isolated from normal, disease free, fresh reduction mammoplasties (n=12; age 18-40, BMI < 30, Fig. 1a). Patient-to-patient variability in organoid generation capacity was evaluated by thoroughly analyzing whole-gel confocal images taken at 5-6 focal planes (Z) and quantifying the number of organoid structures formed. When seeded as single cells all but one patient sample yielded organoids over a 19-day period. Three types of structurally distinct organoids develop in these cultures. (1) The most common type are ductal-lobular structures that consist of branched elongated ducts that terminate with alveoli (60.48% ± 7.96% mean ± STD). (2) Ductal structures with no alveoli are the second most common type (31.17 ± 11.77% mean ± STD), followed by (3) Acinar structures clustered with no visible ducts (8.35 ± 5.06% mean ± STD) (Fig. 1b, c). Significant variability is observed in average organoid formation frequency between patients, (Fig. 1d), and no correlation was found between regeneration capacity and patient age (R^2^=0.007) or BMI (R^2^=0.003). Across the 12 patient samples, the median number of structures formed per 100 seeded cells was 1.775, with a 95% CI between 0.45 and 4.10 (Fig. 1e).

**Fig. 1:**
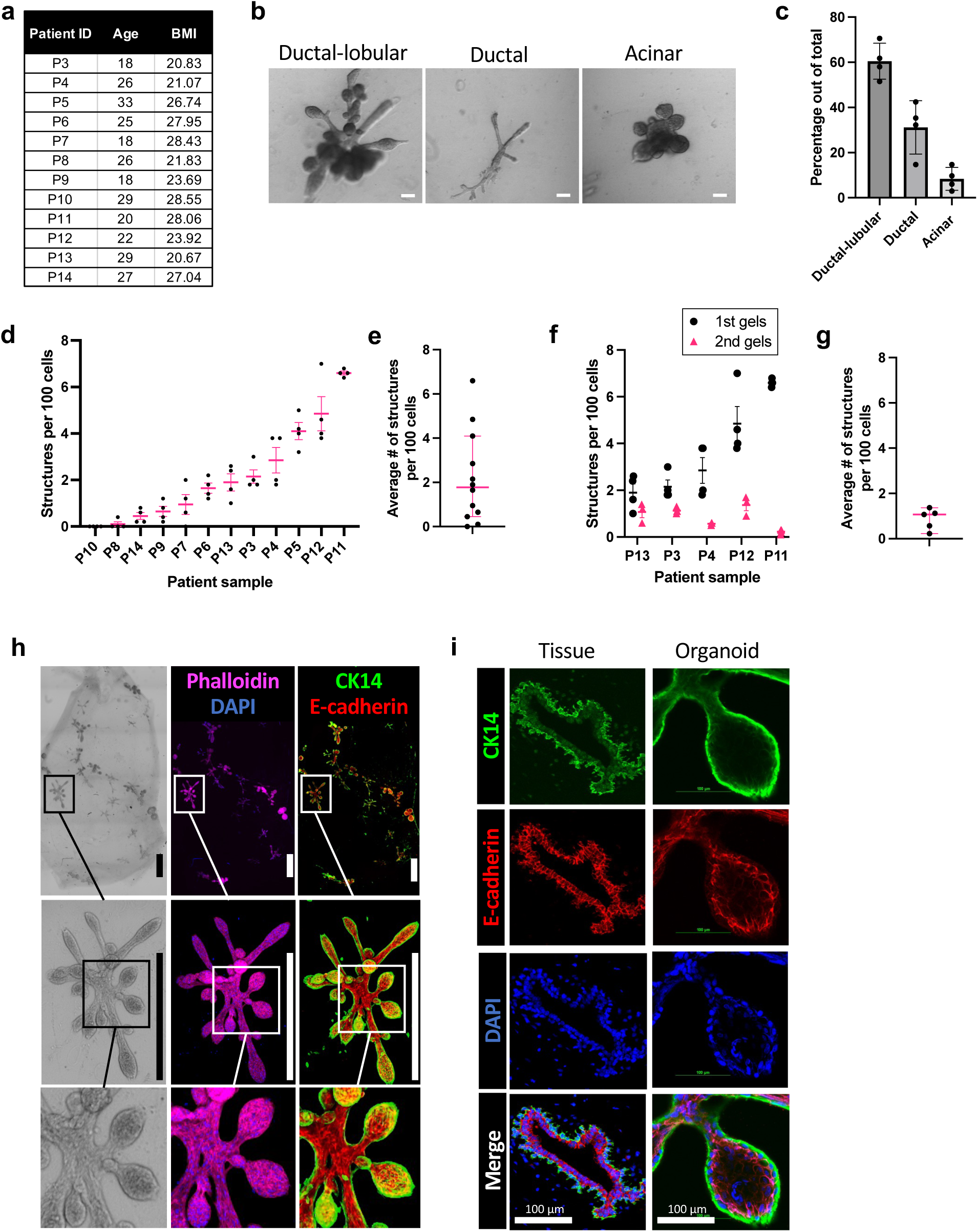
Regenerative and biomimetic capacity of breast organoids. **a**, List of tissue donor patients with age and body mass index (BMI). **b**, Representative bright field images of organoid morphology types. Scale bar = 100 μm. **c,** Proportion of organoid morphologies. Mean ± STD. **d,** Rate of organoid formation per 100 cells from 12 patient samples (n=4 repeats per sample). Mean ± SEM. **e**, Rate of organoid formation across patient samples. Median ± 95% CI. **f**, Rate of primary and secondary organoid formation of 5 patient samples (n=4 repeats per sample). Mean ± SEM. **g**, Rate of secondary organoid formation across 5 patient samples. Median ± 95% CI. **h**, Whole-gel images of bright-field (left panels) and immunofluorescence staining for F-actin (phalloidin, magenta) with DAPI (blue, middle panels), and E-cadherin (red) with CK14 (green, right panels). Scale bars = 1mm. **i**, Organoid (right) and patient tissue (left) immunostaining for CK14 (green), E-cadherin (red) and DAPI (blue). Organoid and tissue are not from same patient. Scale bars = 100 μm.

Secondary organoid formation was performed to evaluate the regenerative capacity of stem/progenitor cells within formed structures. Primary organoids established from five patient samples were dissociated and single cells were re-seeded into a second set of hydrogels (n=4 per sample). The number of secondary organoids that formed by day 17 was quantified. In all five patient samples, we observed a reduction in secondary organoid formation compared to primary organoid formation (Fig. 1f) suggesting a reduction in either stem cell number or regenerative capacity of stem cells. The median number of secondary structures formed per 100 seeded cells was 1.067, with a 95% CI between 0.23 and 1.37 (Fig. 1g). Interestingly, there was greater patient-to-patient variability in primary organoid formation compared to secondary organoid formation, suggesting that once organoids form, they maintain a relatively consistent pool of regenerative cells within them.

Organoids show physiologically accurate architecture, size, and cell patterning of human breast tissue. TDLUs with a network of ducts terminating in alveoli was recapitulated in organoids, with a layered epithelium consisting of luminal cells expressing E-cadherin and basal cells encapsulating them and expressing CK14 (Fig. 1h). Side-by-side comparison of breast tissue sections with organoids show a similar expression pattern of the luminal marker E-Cadherin and the basal markers CK14 (Fig. 1i).

Together these results show that next-generation organoid formation undergoes a series of well-coordinated events that result in the formation of a multi-layered biomimetic breast tissue that accurately resembles complex and heterogenous breast tissue. Since organoids are derived from a single cell, and the resulting complex tissue is comprised of both ducts and acini as well as both basal and luminal cell lineages, this parent–progeny relationship between the precursor cell and the organoid demonstrates they arise from facultative multilineage stem cells.

### Comparative Transcriptional Analysis of Tissue and Organoid Cell Populations

To ascertain if organoids grown in ECM hydrogel mirrors human breast tissue, we performed a side-by-side analysis using previously published single-cell RNA sequencing (scRNA-seq) data on cells obtained directly from normal human breast tissue^14^ as well from traditional organoids grown in Matrigel^15^ and compared them to scRNA-seq we performed on organoids grown in the ECM hydrogel substrate. Briefly, organoids from 5 donors were pooled, dissociated into single cells, and equally represented from each donor for 10x scRNA-seq, where we sequenced 8,276 cells. Using Seurat v5 we integrated the three scRNA-seq datasets using canonical correlation analysis (CCA) integration, and visualized the clustering with t-distributed stochastic neighbor embeddings (t-SNEs). Cell clusters were identified based on canonical gene expression patterns. The integrated data revealed several interesting features (Fig. 2a, b). First, both human tissue and organoids grown in hydrogel ECM was comprised of heterogenous cell types including both epithelial and non-epithelial cells. Not surprisingly, reflecting the greater cellular diversity and intricacy of in vivo tissue, some of the non-epithelial cell types found in human breast tissue included three immune cell populations - T cells, macrophages, and neutrophils as well as vascular cell populations.

**Fig. 2:**
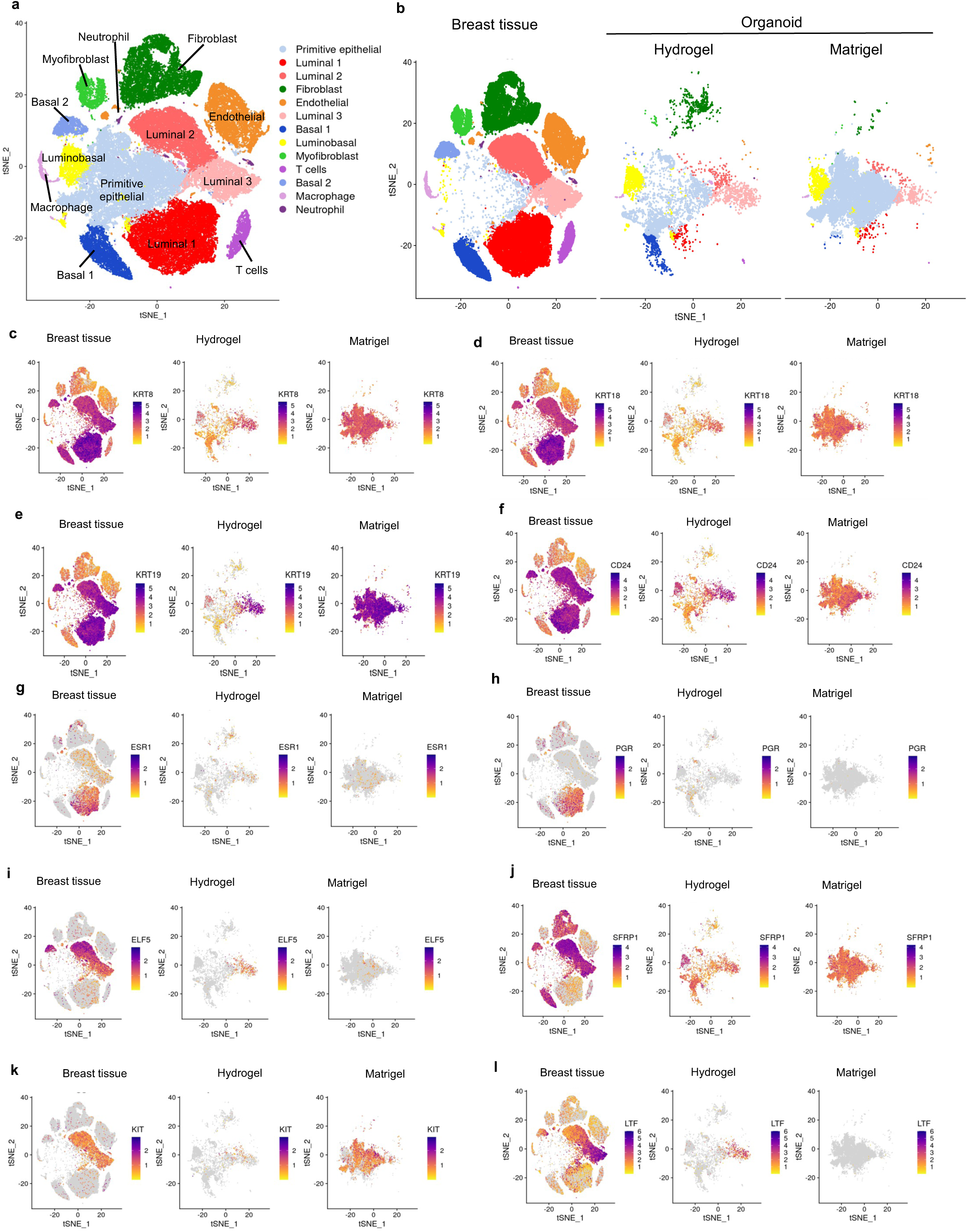
scRNA-seq analysis of breast tissue and organoids grown in ECM hydrogel or in Matrigel. **a-b**, Side-by-side comparison of scRNA-seq clustering of cells derived from healthy human breast tissue and breast tissue organoids. **c-f**, Expression patterns highlighting luminal markers KRT8, KRT18, KRT19, CD24. **g-h,** Expression patterns of hormone receptors ESR1, PGR. **i-k,** Expression pattern of ELF5, SFRP1, and KIT. **l,** Expression pattern of the gene encoding lactoferrin (LTF).

Second, both human breast tissue and organoids grown in hydrogel ECM contained three populations of luminal cells, a luminobasal population, two populations of basal cells, and interestingly a population of primitive epithelial cells (Fig. 2a, b). In addition, a population of mesenchymal/fibroblast cells were also found in both human breast tissue and hydrogel organoids. (Fig. 2a, b). Luminal and basal epithelial cells in hydrogel organoids showed clustering patterns like those of their in vivo counterparts indicating that the cells within the organoids possess transcriptional profiles and traits akin to cells in natural tissues (Fig. 2a, b). All three sub populations of luminal cells were identified: Luminal 1, Luminal 2, and Luminal 3. All three populations expressed the classic luminal epithelial markers KRT8, KRT18 and KRT19, as well as CD24 (Fig. 2c-f). However, Luminal 1 cells expressed the hormone receptors genes ESR1 and PGR (Fig. 2g, h), while Luminal 2 and Luminal 3 cells lacked these genes. Luminal 3 cells expressed ELF5, SFRP1, and KIT linking this population more closely with both luminal progenitor cells and mature luminal cells (Fig. 2i-k). However, Luminal 3 cells also expresses lactoferrin (LTF), a key component of breast milk (Fig. 2l). Luminal 2 cells in contrast, expressed genes associated with both basal and luminal markers, which could be a luminal-basal population (Fig. 2c-f, Supplemental Fig. 1a-c). Basal cells on the other hand expressed the classic basal epithelial markers KRT5, KRT14, and KRT17 (Supplemental Fig. 1a-c).

Third, not surprisingly, out of the populations shared by the organoid models and breast tissue, organoids grown in Matrigel only contain ∼1% luminal cells and instead contain luminobasal cells (7.3%) with the vast majority of organoids being a uniform hybrid basal cell population characterized by the expression of luminal, basal and EMT genes simultaneously (88.33%) (Supplemental Fig. 1d). This population is also enriched for proliferative genes (KI67, PLK1, MYBL2, TOP2A), which might be akin to cap or body cells of the developing terminal end bud, or they may be organoid-specific adaptation of basal cells to the culture conditions (Supplemental Fig. 1e-h). This population lacks a direct counterpart in vivo, and is only 48.67% in the hydrogel. Given that Matrigel mimics the basement membrane and influences the differentiation and proliferation of breast epithelial cells, this ECM scaffold seems to favor the development of an epithelial cell population that might not accurately replicate the cellular diversity of natural breast tissue.

Similar to human breast tissue, organoids grown in hydrogels contain a prominent luminal and luminobasal population (34.35%) as well as a basal population (7.94%) and a prominent population of primitive epithelial cells (48.67%) that is not typically observed in native tissue and may not have a direct counterpart in the adult human breast.

Fourth, both human breast tissue and organoids grown in hydrogels also have a moderate presence of mesenchymal/fibroblast cells making up 20% of the cells in human breast tissue and nearly 9% in hydrogel organoids. In contrast, organoids grown in Matrigel did not exhibit this mesenchymal population. This non-epithelial mesenchymal population within the organoid cultures corresponded to stromal fibroblasts found in vivo (Fig 2a, b) and expressed the stromal markers PDGFRB, COL1A2, COL6A3, THY1, and ZEB1 (Supplemental Fig 2a-e).

These results demonstrate that hydrogel organoids recreate a significant portion of the cellular diversity of actual breast tissue including the varied functionalities of luminal cells as well as mesenchymal cells. The scRNA-seq results highlight the hydrogel organoid’s ability to mimic distinct cell populations observed in natural breast tissue, with the luminal and basal cells particularly resembling their in vivo counterparts. However, there are differences in the prevalence of Luminal 1 and Luminal 3 populations relative to Luminal 2 in organoids compared to that of human breast tissue. This discrepancy could be due to differences in the later stage hormone-driven developmental changes reflected in vivo that is not present in vitro. For example, exposure to puberty and pregnancy hormones drives luminal cell expansion and differentiation, as do the hormonal fluctuations of the menstrual cycle, which influence cell type proportions and cellular composition of breast tissue ^16^. Neither of these events have yet been recapitulated in vitro nor have the tissues been grown for longer than 28 days. Future studies will be needed to explore long-term cultivation and hormonal treatments mimicking puberty, pregnancy, or lactation stages to determine if the cell profiles of organoids more closely reflect in vivo conditions. Together, this affirms our organoid’s capacity as a biologically relevant model for studying breast tissue biology and associated disorders.

### Non-canonical cell dynamics, ductal morphogenesis, and epithelial folding during organoid formation revealed by high-throughput, long-term live confocal imaging

We modified the organotypic culture methodology to enable high throughput and high-speed live-imaging confocal microscopy (Fig. 3a). We scaled down the hydrogel volume from 200 to 20 μl gels to accommodate a 96-well format. Patient-derived primary breast epithelial cells were labeled with a cell tracking dye (Cytopainter Green) to distinguish them from cell debris on the day of seeding. This step was important to improve the visualization and ensure the tracking of live primary cells, which are difficult to distinguish morphologically at this early stage. Labeled cells were seeded at a density of 100 cells per hydrogel and cultured for 2.5-3 weeks. Cultures were kept in a stage-top incubator that maintained controlled temperature, CO_2_ levels, and humidity, all within an enclosed dark chamber attached to a confocal microscope. We live tracked 111 locations on the plate, capturing 9 Z planes spanning 160 μm every 30-45 minutes over the 18-21 day period. This methodology allowed for unprecedented visual documentation of organoid formation in real time.

**Fig. 3:**
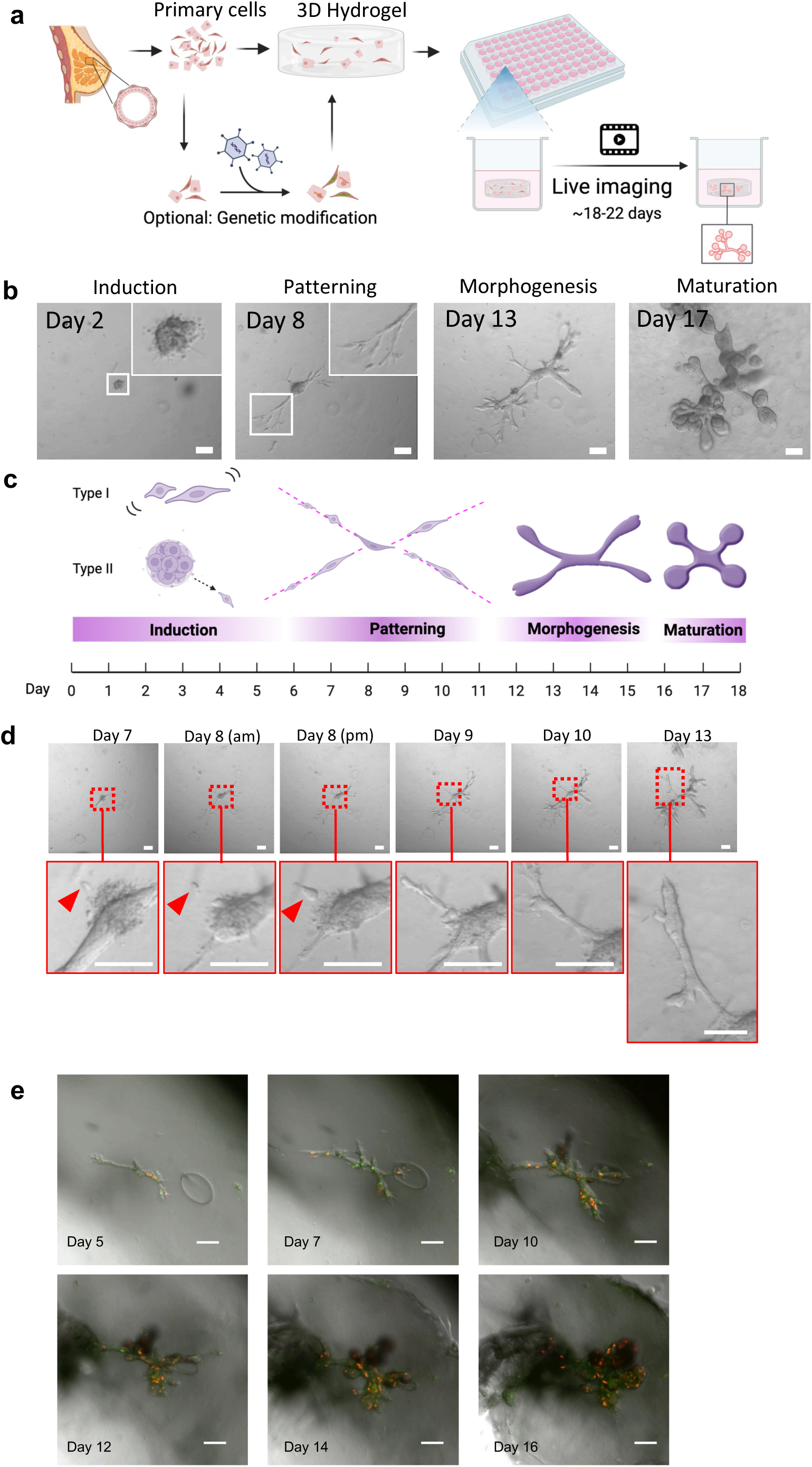
Live Imaging of Cellular Dynamics in Breast Organoid Development. **a**, Schematic representation of the high-throughput confocal live imaging setup for observing human breast organogenesis and maturation in the organoid model. The diagram details the experimental design and methodology. **b**, Panel of selected frames from Supplemental Video 1, demonstrating the four phases of organoid development, with Type I induction. Scale bar = 100 μm. **c,** Schematic representation and timeline of the 4 phases of organoid development, including Type I and Type II induction. **d**, Panel of selected frames from Supplemental Video 1, showing type II induction culminating with cell (red arrowheads) detaching and moving away, branch subsequently grows in the direction of the preceding cell. Scale bar = 100 μm. **e,** Panel of images displaying the development of a breast organoid, starting from a single cell and infected with a FUCCI vector to display changes in the cell cycle, between days 5-18. Cells displaying red are in G1 phase, while cells fluorescing green are actively dividing and are in S, G2, or M phases. Scale Bar = 100 μm.

We observed that organoid formation occurs in 4 distinct phases: Induction, Patterning, Morphogenesis & Maturation spanning approximately 18 days. The full process of organoid development, encompassing all four phases, is illustrated in Fig. 3b, c and Supplemental Videos 1-6. Inductive interactions, especially between the epithelium and the ECM is crucial for organ initiation ^17,18^ and can be seen in real-time in this model. We observed organoid induction as a distinct preliminary stage, taking place from days 0 to 8. During the Induction Phase, cells exhibit minimal movement and largely stay stationary, with the initiation of cell proliferation. We observed two distinct induction types: Type I, the predominant type that starts with isolated, dispersed, spindle-shaped cells (Fig. 3b, c and Supplemental Videos 2-5) and Type II, that begins with close-packed cells in a spheroid from which individual elongated, spindle cells emerge. The ejection of such cells from a spheroid colony initiates the next phase - Patterning. (Fig. 3d and Supplemental Videos 1, 4).

Over the next ∼10 days, colonies begin to pattern into rudimentary branches in a dynamic process of cell trafficking where individual cells migrate back and forth along the axis of the extending branch. Single cell tracking of this dynamic could be readily observed when primary cells were infected, prior to embedding in the hydrogel with a lentiviral vector expressing the fluorophore Venus (LeGO-V2, Addgene plasmid #27340 ^19^).

Although Venus expression varies during the patterning phase, patterning evolves as motile cells scatter, setting a preliminary design for the branching pattern of the organoid. A single Venus-positive cell can be observed moving from the tip of the invasion front to the organoid’s main body and then returning to the front (Supplemental Video 4). This initial design is later populated by cells that form the branched structure during the subsequent Morphogenesis phase. This process of ductal patterning contrasts with the classic branching process of ductal elongation that proceeds through steady invasion of a fully formed bud spearheaded by leader cells^20,21^.

Morphogenesis is marked when cells begin to both proliferate rapidly and rearrange themselves to form complex three-dimensional shapes, including tubes, buds, and cavities. By day 16-19, organoids continue to grow and start to mature and develop buds. During the Maturation Phase, the tissue assumes the characteristic structure of a TDLU, comprising round alveoli that are closely grouped and interconnected (Fig 3b, c). Notably, fluorescent labeling revealed substantial coordinated movements and rearrangements among cells within the evolving TDLU. This coordination is evident in the spiral rotation observed in the terminal acini, as well as in the longitudinal migration and stretching along the ducts (Supplemental Videos 1, 4).

Single cells were also infected, prior to embedding in the hydrogel, with a lentiviral vector expressing the cell cycle reporter FastFUCCI ^22^ to gain a deeper insight into proliferation dynamics during organoid formation. Cells transduced with FastFUCCI fluoresce different colors as they transition through the cell cycle. This enables real-time visualization of cells actively cycling in S, G2 and M phase of the cell cycle, which fluoresce in green (mAzami-Green), while following cell division, in the early G1 phase, cells display no fluorescence^22^. However, as they progress through the G1 phase, orange fluorescence (mKO2) gradually intensifies, allowing G1 cells to be easily tracked within the expanding organoid. FastFUCCI labeling revealed a diverse mix of orange, green, and non-fluorescent cells throughout organoid formation. Cells quickly move through the G1, S, G2, and M phases of the cell cycle. Early during organoid formation cells are mostly negative or green suggesting little time in G1. However, as development progresses an accumulation of G1 orange cells can be observed, which are dynamically moving and migrating throughout the growing TDLU (Fig. 3e, Supplemental Video 7). By the maturation phase, orange G1 cells dynamically reposition themselves along the ducts, and within the acini and exhibit considerable motility.

### Cell tracking of organoid formation reveals diverse movement and growth patterns during organogenesis

We designed a bioimaging analysis pipeline to quantify organoid growth during organogenesis by measuring changes in volume expansion and surface area over time. This analysis was conducted using a dataset comprising 11 videos, tracking the development of 11 distinct organoids (Supplemental Videos 11-19). Growth was measured throughout organoid development, and the final organoid size was recorded (Figure 4a-e, Supplemental Figure S3-S4). We observed that some organoids exhibited the predictable Gompertz growth pattern of an initial lag, followed by exponential growth, and then transitioning through a deceleration phase before reaching a plateau as they mature (organoids 1, 2, 6; R² >0.9), while others exhibited a linear model of growth that was constant over time (organoids 5, 7; Log R² >0.9) (Supplemental Figure S5-S6). Other organoids did not fit either model of growth. In addition to modeling the different growth trajectories for each organoid, we also measured their growth rates. Organoid growth rates varied with some rapidly growing (e.g., organoid 6, slope = 2.662), and others exhibiting moderate or slower growth rates (e.g., organoid 9, slope = 0.2437). We found a strong positive correlation between the log-linear growth rate and its R² values (r = 0.811, n = 12, p < 0.05), and a moderate negative correlation between the Gompertz growth rate and its R² values (r = -0.566, n = 10, p < 0.05). This indicates that organoids that grow faster tend to follow the exponential growth model more closely, but faster-growing organoids may not always grow in the smooth, predictable Gompertzian pattern that was expected. This could be due to the rapid transitions through the lag, growth, and plateau phases.

**Fig. 4:**
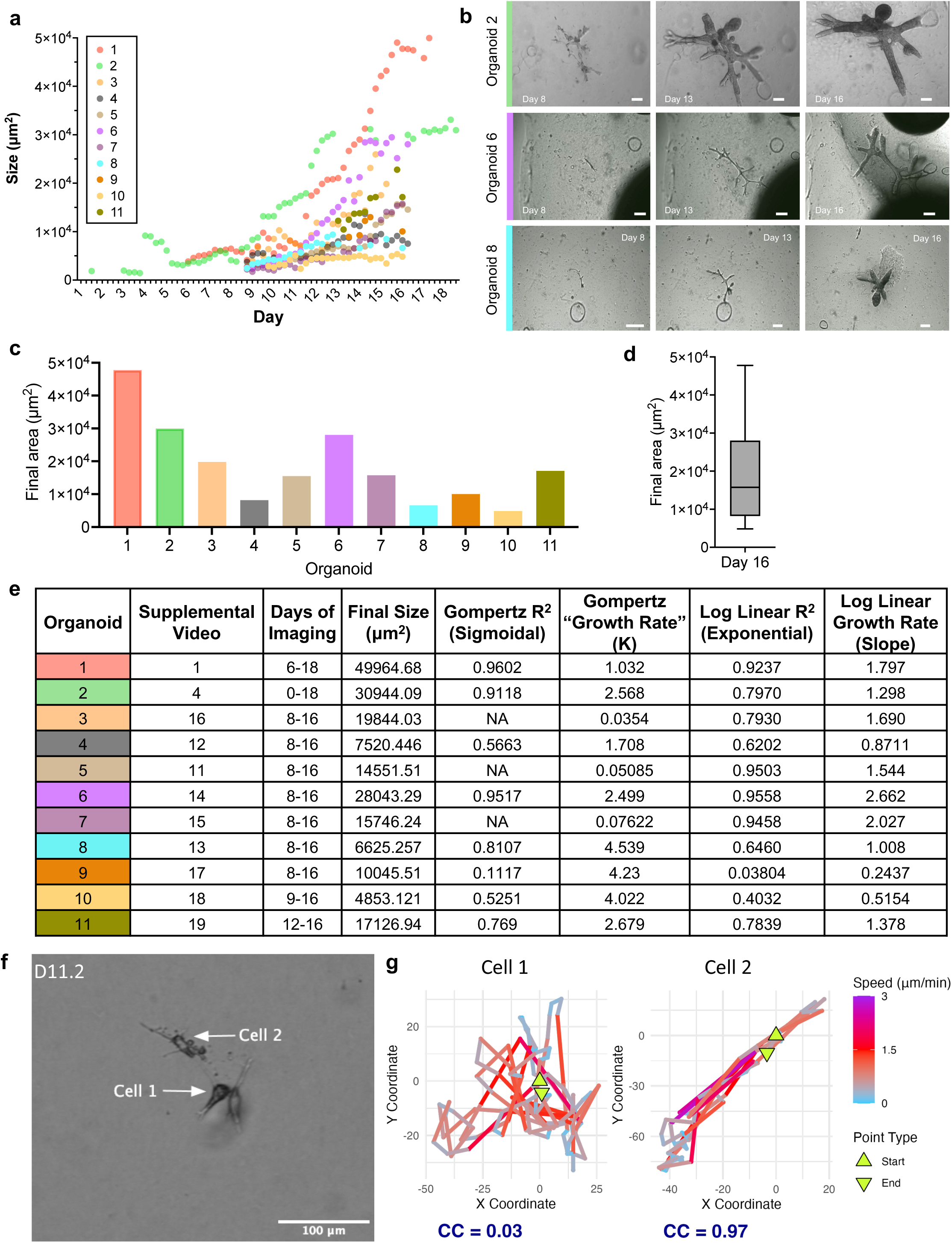
Tracking organoid growth and cell dynamics during organoid development. a, Scatter plot depicting the area of 11 organoids during their development. Area measuremtnes were averaged every 6 hours. **b,** Representative brightfield images of organoids 2, 6, and 8 from the time-course plot in (a), (color-coded corresponding to the scatter plot) on days 8, 13, and 16, highlighting the morphological changes and growth progression of individual organoids over time. **c,** Final area of each organoid at the end of the observation period (day 16). **d,** Box plot presenting median and quartile data of for a comparable distribution of the final organoid areas. **e,** Table displaying the details of organoids used for quantification of growth. The table includes the following information for each organoid: the corresponding Supplemental Video, the imaging timeframe, final organoid size on day 16, linear regression R^2^ values and slopes for Gompertz and log-transformed data. Each row is color coded to match (a)-(c). **f,** Single frame image from live imaging video. Arrows indicate two tracked cells, labeled Cell 1 and Cell 2. Scale bar = 100 µm. **g,** Trajectory plots of Cell 1 and Cell 2. The color gradient indicates the speed of the cells. Triangles denote the starting and ending points of the tracks. The correlation coefficient (CC) between the X and Y coordinate values is denoted for each cell track.

We also assessed whether there was any correlation between growth rates and their final size or shape. Indeed, there is a moderate positive relationship between the final size of the organoids and their exponential growth rates (r = 0.575, n = 12, p < 0.05), consistent with the notion that organoids with higher growth rates tend to achieve larger final sizes. However, the correlation is not strong, indicating that while growth rate is an important factor, it is not the sole determinant of the final size of organoids. Qualitatively, it appears that the potential to grow to a certain size is influenced by its shape, with branched organoids generally exhibiting larger growth compared to spherical and irregular shaped organoids. Further quantitative analysis is required to confirm these findings.

Taken together these interesting differences in organoid growth patterns and rates could be due to the variability in stem cells that initiated organogenesis, differences in cellular metabolism, or even slight differences in niche or cellular microenvironment within each growing organoid. Studying these differences might provide insights into cellular behaviors during complex tissue formation.

We next tracked individual cells in each frame of a time-lapse video, during induction and patterning when organoid growth was at its lowest rate. We were able to track individual cell movements, measuring cell velocity, directionality, and displacement to gain a deeper understanding during the initial stages of organoid development. This revealed cell velocity and patterns of movement across different cells during induction and patterning, and enabled the calculation of a correlation coefficient (CC) between the x and y coordinates of each cell trajectory, (Fig. 4f, g, Supplemental Fig. S7). Some cells exhibited linear or smooth curves with CC values closer to 1, while others displayed jagged or erratic path patterns with CC values closer to zero, that indicate random or exploratory movement (Supplemental Fig. S7e, f) . In these cases, the cell may frequently change directions, suggesting response to complex stimuli or interaction with the environment. Looping or circular paths had varying CC values depending on the consistency of the looping direction (Supplemental Fig. S7g, h). Forward and backward movements along similar lines was defined as CC closer to 1. Random movement was defined as CC values closer to zero and plots where the cell path is highly irregular, with frequent direction changes without progressing far from the origin. Together, these diverse behaviors are indicative of different environmental cues, intrinsic cell properties, and stages of the cell cycle. In some cases, cells with CC close to 1 exhibited reversing movement pattern where the cell moves back-and-forth even returning to their starting point, or exhibited largely linear, targeted movements with directed trajectory at high speed (Figure 4g). This diversity in cell movement during induction and patterning may indicate different functional roles for the cells in the organoid’s structural development. In addition, this variability in movement patterns during the early stages of organogenesis likely reflect an inherent flexibility in the developmental program of the organoid. Further study will be required to elucidate the importance in the cell behaviors and movements during organogenesis and organoid development.

### Mapping the emergence of and characterizing organoid mesenchyme during organogenesis

Time-lapse imaging of organoid development revealed the unexpected formation of a highly motile population of mesenchymal-like cells that scatter away from the edges of the growing organoids (border cells) and rapidly proliferate (Supplemental Videos 2, 5, 8-10). This population of rapidly proliferating mesenchymal-like cells can be seen dynamically interacting with the collagen matrix and actively contracting and manipulating the collagen matrix, causing the hydrogel to bend and fold. They cover the hydrogel’s outer surface, moving in a synchronized, wave-like sinusoidal pattern. As a result, the hydrogel eventually condenses into a sphere, with epithelial TDLU organoids encased inside, and enveloped by a layer of flat, spindle-shaped cells on its exterior surface.

The very active, dynamic, and multilineage nature of organoid formation is reminiscent of events observed during embryogenesis. During embryogenesis, the early ectoderm is the precursor to both the stromal and epithelial components of the breast. Interestingly, live imaging videos reveal that highly motile mesenchymal cells originate from the same cells from which the organoid arises suggesting a common precursor (Supplemental Video 5). During embryogenesis, the expression of epithelial markers such as CK14 and E-Cadherin begin during the process of epithelialization ^23–26^. Notably, we found that during induction, neither CK14 nor E-cadherin are expressed in the single cells that have not yet formed an organoid (Fig. 5b). However, by day 10 day, the epithelial marker CK14 is expressed when basal epithelial cells appear, consistent with commitment to the epithelial lineage. E-cadherin, which is crucial for epithelial cell-cell adhesion is expressed when luminal cells appear within the inner layer of the thickening branches by day 15. This localization of E-Cadherin during morphogenesis is most predominant at locations within structures that will become the alveoli (Fig. 5b).

**Fig. 5:**
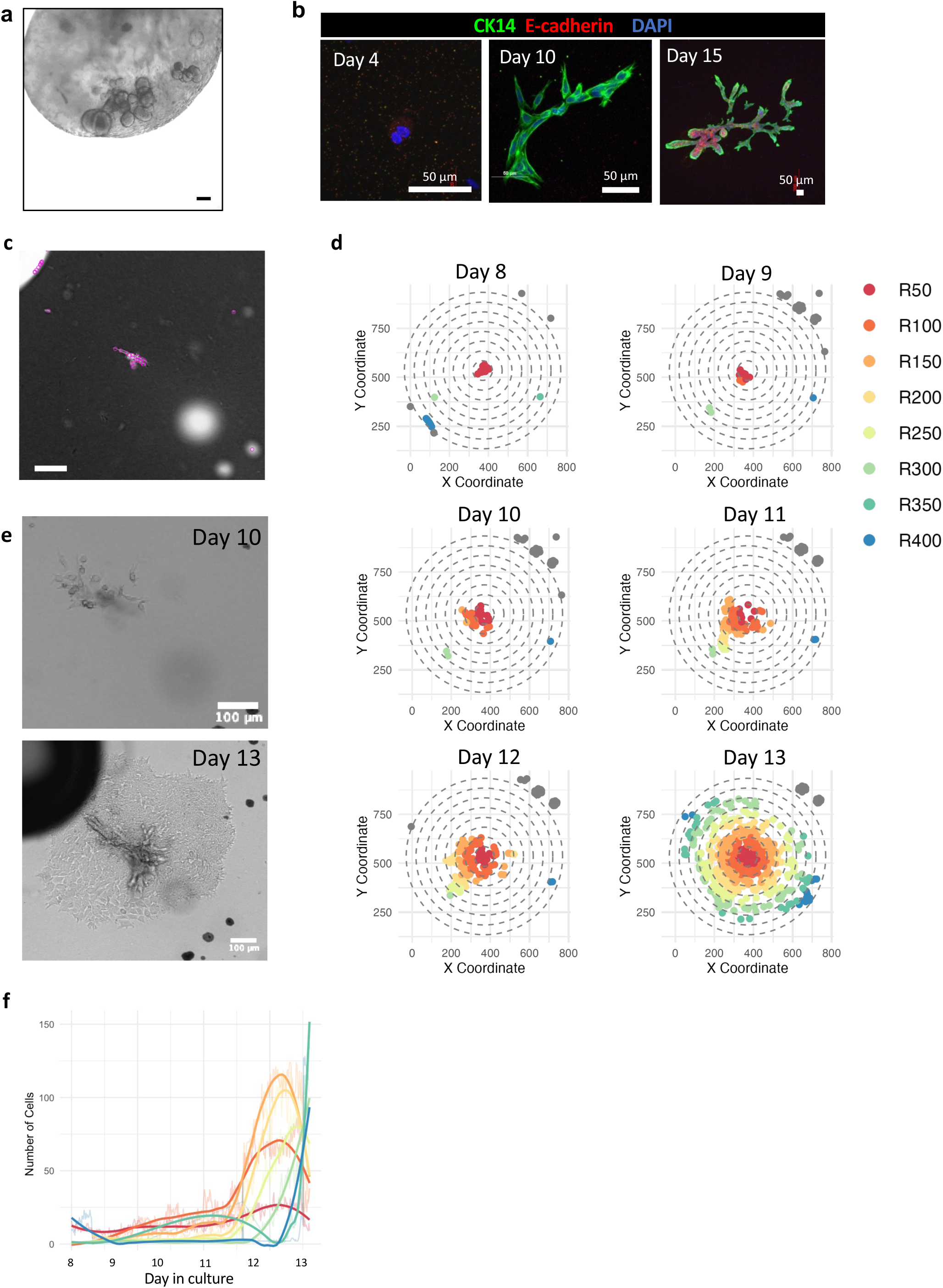
Tracking cell populations demonstrates radial expansion from organoid origin point. **a,** Frame from Supplemental Video 5: Mature organoid producing the classic morphology of a terminal ductal-lobular unit embedded within compacted mesenchyme. Scale bar = 100 μm. **b,** Immunostaining of organoids at different stages of development. CK14 in green, E-cadherin in red and DAPI in blue. Scale bars = 50 μm. **c,** Representative frame from live imaging video (inverted bright field). Purple circles mark spots detected by the cell tracking software (TrackMate). Scale bar = 100 μm. **d,** Plots showing the spatial distribution of cells at various distances from the center over time. Each plot represents a different time point. Cells are color coded corresponding to their distances from the center, as indicated in the legend on the right (e.g., R100 indicates distance of 100 μm from the center). **e,** Bright field images from live imaging videos showing organoid development on days 10 (top) and 13 (bottom). Scale bar = 100 μm. **f,** Graph depicting the number of cells over time for each distance category from the center. Colored line corresponds to distance categories, matching the plots in ‘b’. Raw data lines are shown in the background and smoothed lines in the foreground.

We applied automated tracking software with deep learning-based segmentation algorithms and Laplacian of Gaussian (LoG) filter for object detection to automatically detect individual cells emerging from organoids in each frame of a time-lapse video to assess the origin of the mesenchyme. If mesenchymal cells emerge from the organoid rather than from contaminated stromal cells, a single central gradient of cell density of growth would be observed. Contaminated stromal cells seeded within the gels would exhibit growth patterns such as random distribution, multiple clusters, or edge-biased growth, depending on the initial seeding location and cell behavior.

We analyzed two live imaging videos (Supplemental Videos 5, 8) and observed a similar pattern of mesenchyme emergence: a central concentration of cells that distribute radially as time progresses, actively proliferating and spreading from the organoid center. We did not find evidence of a uniform or multifocal proliferation of cells away from organoids (Fig. 5c-d, Supplemental Fig. S8, Supplemental Videos 5. 8). On day 8 of organogenesis, cells are concentrated near the center (red and orange spots) and there are a few scattered cells at the periphery (blue and green). By day 10 and 11 there is an increased concentration of cells near the center and the region of dense cell concentration is expanding radially. By day 12 and 13, there is further expansion of the dense cell region which becomes the organoid parenchyma, and the mesenchymal cell distribution becomes more spread out from the center. The increase in the area covered by different colors suggests an active expansion and dispersion of cells from the initial seeding point. The continuous presence of cells at different radii over time also implies sustained proliferation.

During embryogenesis cells positioned at the boundary of different tissue regions also often undergo epithelial-mesenchymal transition (EMT), such as the case during neurulation. Border cells at the edges of the neural plate contribute to the bending of the neural plate and the closure of the neural tube. Given the similarities between neural plate closure and the folding and bending of the hydrogels, we examined whether the border cells and mesenchymal-like cells exhibited features of EMT. Immunohistochemistry (IHC) analysis revealed a mixed population of activated mesenchymal-like states characterized by a spectrum of EMT stages, from fully transitioned mesenchymal cells to those still undergoing EMT. Differential expression of EMT markers (CK14, Zeb1, Snail) alongside mesenchymal markers (S100A4, FAP, FSP, Vimentin, CD90/Thy1) that lacked epithelial makers could be seen within the organoid mesenchyme (Fig. 6a-f, Supplemental Fig. S9).

**Fig. 6:**
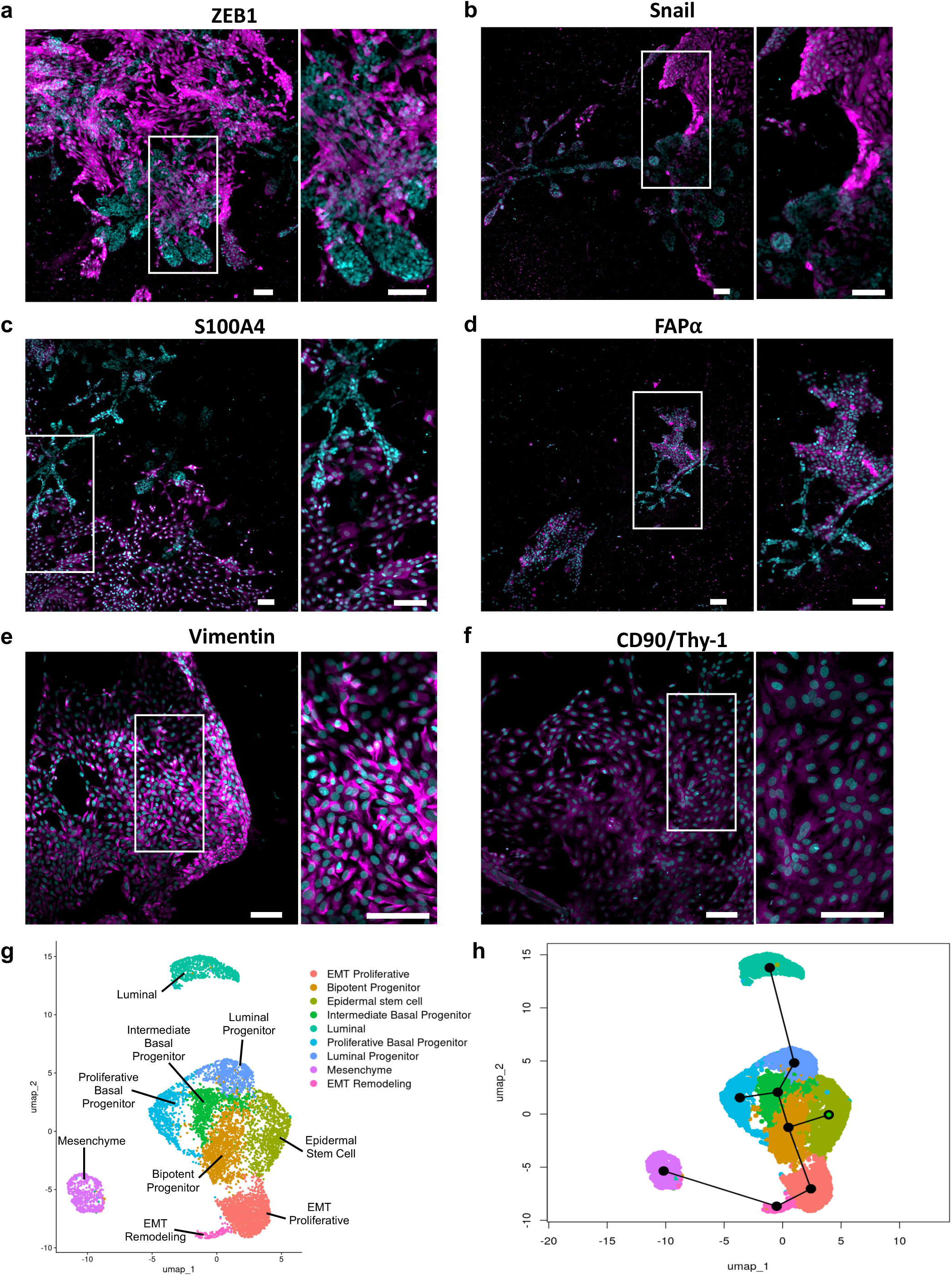
Expression of epithelial and mesenchymal markers in maturing organoids. **a-f,** Immunostaining of mesenchymal-like cells that surround epithelial organoids: ZEB1 (a), Snail (b) S100A4 (c) Fibroblast Activation Protein alpha (FAPα, d), Vimentin (e), CD90/Thy-1 (f). Magenta: protein markers; cyan: Hoechst nucelar staining. Right images in each panel represent higher magnification images of the regions indicated by the white boxes in left image. Scale bars = 100 µm. **g,** UMAP of scRNA-seq of hydrogel organoid cells colored by cell type. **h,** Pseudotime analysis of hydrogel organoid clusters. The lines represent the three lineage trajectories with the origin highlighted in green.

To further identify the origin of the mesenchymal population, we analyzed the scRNA-seq data from organoids cultured in ECM hydrogels using Seurat v5, and performed pseuditime analysis using Slingshot ^27^ (Fig. 6g, h). Cells were analyzed using unsupervised graph-based clustering and were visualized using Uniform Manifold Approximation and Projection (UMAPs). We identified 9 cell clusters, and these were identified as specific cell types based on differential gene expression of canonical cell markers. The identified cell clusters include epidermal stem cells, intermediate basal progenitors, luminal cells, luminal progenitor cells, as well as a distinct separate mesenchyme population. Bridging the epithelial and mesenchyme populations are proliferative and remodeling EMT clusters (Fig 6g).

The luminal cluster expresses markers expected in mature luminal cell types such as EPCAM, KRT8, KRT18, KRT7, CD24, KRT19, as well as LTF (Supplemental Fig. S10a-g) . All of the basal cell clusters express the expected markers KRT14 and KRT17 (Supplemental Fig. S10 h, i), but the proliferative progenitor cluster expressed increased proliferative markers MKI67 and TOP2A (Supplementary Fig. S10 j, k). The epidermal stem cell population was identified based on its overlapping expression signature with known markers for epidermal differentiation and keratinocyte function, such as KRT15, KRT6C, DSG1, DSC2, ZNF750, SPINK5, EVPL, and S100P (Supplemental Fig. S10 l-s). The mesenchyme population expresses high levels of MME, THY1, ZEB1, FAP, and S100A4 (Supplemental Fig. S10 t-x). The EMT proliferative and remodeling clusters have overlap between epithelial markers and mesenchymal markers, including Vimentin, CDH3, CDH1, and Slug (Supplemental Fig S10 y-ab). Both by IHC and scRNA-seq, basal epithelial cells express EMT markers such as CK14, Vimentin, and Slug (Fig. 1h, i; Fig. 5b; Fig. 6b; Supplemental Fig. S9a, c; Supplemental Fig. S10 h, y, ab, but so do many of the mesenchymal cells as well. This overlap in EMT markers in two very different tissue compartments in the organoid cultures, reinforces the notion that the stromal cells are a heterogenous mixture of EMT and mesenchymal populations that contribute to distinct tissue structures.

Pseudotime analysis done using Slingshot reinforces the lineage distinctions of the cell clusters, with three separate linages identified (Fig. 6h). Based on the hypothesis that the primitive epidermal stem cells are the initiating cells within the hydrogel grown organoids, we set the pseudotime 0 at this cluster. The initial branch point of the lineages is at the bipotent progenitor cluster, with one lineage leading towards the EMT and mesenchyme clusters, while the other two lineages diverge at the intermediate basal progenitor cluster. The second lineage terminates at the proliferative basal progenitor cluster, with the final lineage terminating at the luminal cluster. This supports that in the organoid model the mesenchymal cells and the epithelial cells arise from the same cell population.

### Impact of menopause on organoid formation

We next explored the potential of this model to investigate how menopause affects breast stem cell activity during organoid formation. Two key hormones, estrogen and progesterone, are pivotal for breast development and stem cell function ^28,29^, and their decline following menopause is linked to breast tissue atrophy ^30,31^. However, due to limited model systems, the functional implications of these hormonal changes on stem cell function and breast tissue regeneration remain unexplored.

We seeded cells from breast tissue collected from breast reduction surgeries of pre- and post-menopausal patients (n=5 patients for each group). Organoids were cultured for 14 days, and structure formation was scored. Compared to pre-menopausal patient samples, cells from post-menopausal patients had a statistically significant 2.91-fold reduction in the rate of organoid formation (1.24 ± 0.72 and 0.43 ± 0.24 (mean ± SD), respectively) (Fig. 7a). These results suggest a loss of facultative stem cell activity in post-menopausal breast tissue. TDLU-like ductal-lobular structures were only observed in pre-menopausal samples and not in post-menopausal ones (Fig. 7b).

**Fig. 7:**
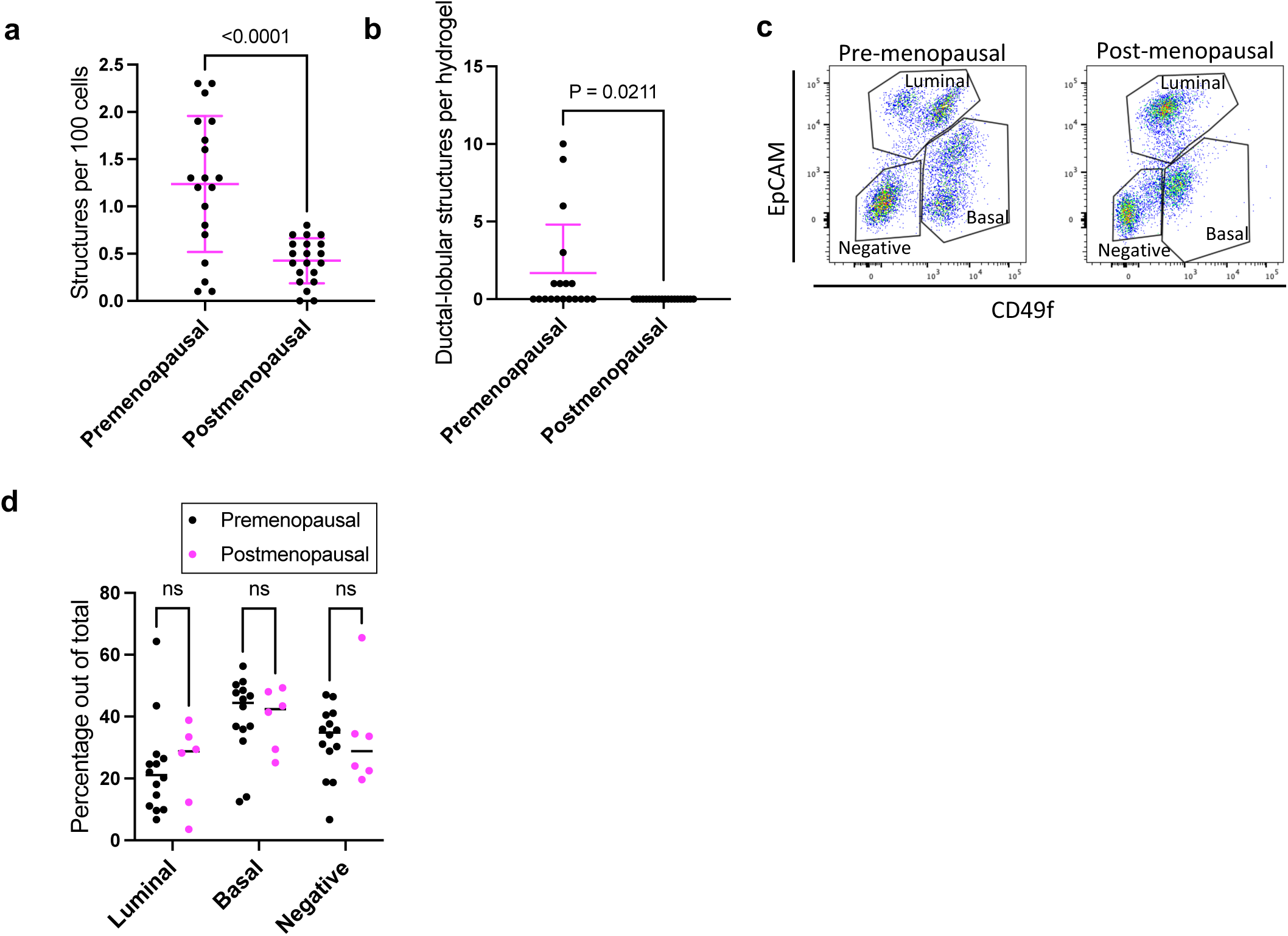
Human Breast Tissue Regeneration and Morphology in a Hormone Responsive Model. **a**, Comparison of the rate of organoid formation per 100 cells between breast cells from pre-menopausal and post-menopausal patients (n=5 patients, 4 gels per patient). Mean ± SD. Statistical comparison: unpaired t-test. **b,** Number of ductal-lobular organoids per hydrogel formed from pre-menopausal and post-menopausal patient samples. Mean ± SD. Statistical comparison: unpaired t-test. **c,** Representative plots of flow cytometry analysis for luminal (EpCAM) and basal (CD49f) epithelial markers. Gating of basal, luminal and non-epithelial (Negative) populations in pre-menopausal and post-menopausal samples. **d,** Percentage of epithelial (luminal and basal) and non-epithelial (Negative) cell populations among pre-menopausal and post-menopausal patient samples. (one way ANOVA with Sidak’s multiple comparisons test).

Using flow cytometry analysis based on detection of luminal (EpCAM) and basal (CD49f) epithelial markers, we compared the epithelial (luminal and basal) and non-epithelial populations among pre-menopausal (n=14) and post-menopausal (n=6) samples to determine whether reduced organoid formation from post-menopausal breast tissue was due to differences in the proportion or types of epithelial cell types seeded (Fig. 7c, d). Three cell populations were identified: luminal (EpCAM^high^), basal (EpCAM^neg/low^/CD49f^pos^), and non-epithelial (Negative; EpCAM^neg^/CD49f^neg^). While the percentage of population did not statistically differ between pre- and post-menopausal samples (Fig. 7d), notably, in most pre-menopausal samples we observed two sub-populations of basal cells, categorized as EpCAM^neg^ and EpCAM^low^, as well as two sub-populations of luminal cells, identified as CD49f^neg^ and CD49f^pos^ (Fig. 7b). In contrast, post-menopausal samples did not have these sub-populations, and showcased a singular basal cell type, EpCAM^neg^, and a single luminal type, CD49f^low^. Together, these results indicate that post-menopausal involution results in decreased cell regeneration capacity and effects on epithelial cell diversity.

Organoids grown in hydrogels express the estrogen receptor (ERα), and respond to estrogen (E) treatment (Supplemental Fig. 11a). We tested whether treatment with estrogen and progesterone (E+P) might affect the regenerative capacity organoids. Hydrogels seeded with cells derived from premenopausal or postmenopausal patients (n=5 for each group) were cultured for 14 days in the presence of estrogen (1 ng/mL) and progesterone (1 ng/mL). We analyzed the effect of hormone treatment on total number of organoids and organoid morphology, and did not find significant differences between the control and treated groups under these experimental conditions (Supplemental Fig. S11b, c).

## Discussion

The inherent ability of cells to organize themselves is a cornerstone of organoid technology. Yet tissue development in vivo does not proceed via spontaneous self-assembly but is driven by directed organogenesis programs. In this study, we have advanced the current paradigm of organoid technology by pioneering a method that promotes directed tissue organogenesis as opposed to spontaneous self-assembly seen in traditional organoid models. This nuanced approach enables the production of organoids that more faithfully replicate in vivo tissues. For the breast, we found that these organoids are indistinguishable from human breast tissue on an anatomical, cellular, and molecular level. This is invaluable for studying cellular processes, interactions, and signaling pathways that drive development, aging, tissue repair, and even cancer. Having a more realistic model which includes both epithelial and stromal cells allows for the study of cancer formation, progression, and stromal-epithelial interactions that are not possible with current 3D models. Having realistic organoid models also enables drug testing in conditions that closely resemble human tissues. This increases the chances that drugs effective in organoid models will also be effective in patients. Moreover, organoids can easily be grown from individual patients, allowing for personalized medicine approaches where drug responses can be predicted for a particular individual. Finally, using this organoid model and tracking method, it will now be possible with the aid of various molecular and genetic labeling tools, to observe how cells divide, migrate, transdifferentiate, and interact with each other and with the extracellular matrix to form tissues in real time.

In addition to the technological advancement, this new method has also provided several additional insights. First, we observed that breast organoid formation reactivates a latent developmental program emphasizing induction, patterning, morphogenesis, and differentiation from single cells. Traditional breast 3D organoid models that spontaneously self-assemble in Matrigel or Type I Collagen mimic only rudimentary aspects of development. Cells adhere to specific neighboring cells, establish polarity, form lumen, and create multicellular structures, by leveraging their intrinsic properties ^32^. The physical environment of the hydrogels however, combined with the live imagining has allowed for the visualization of epithelial induction and patterning in an unprecedented way. We observed two types of induction both of which were foundational phases of organoid development but exhibited different behaviors. In Type II induction, the static behavior of cells appeared to be primarily geared towards preparation for further growth and differentiation that culminated to the ejection of cells from the static colony. In Type I, the initiation of cell proliferation and migration suggests that the cells are preparing for a phase of rapid growth or expansion. Further studies are needed to understand the cellular, molecular, and environmental basis of these types of induction.

We also observed that ductal branching during organoid formation arises from coordinated cellular proliferation and spatial patterning akin to mammary ridge formation during embryonic development, followed by the proliferation of ductal cells and differentiation into lobular and alveolar structures. This developmental process is driven at the tips of the growing ducts by leader cells ^10,33^.

While the participation of leader cells and group cell movement in branching morphogenesis is well documented, the patterning phenomenon observed here, where cells are entirely detached from the branch, and precedes branching by days, has not been reported. Additional studies are needed to further understand this newly identified process.

This breast organoid system recreates certain aspects of breast tissue development but does so by employing mechanisms that are distinct from those observed in traditional in vivo development. In vivo, the first morphologic evidence of mammary gland development begins with ectodermal thickenings, termed mammary ridges or milk lines, that develop on the ventral surface of the embryo. The remnant of the mammary ridge ectoderm begins to proliferate and forms the primary mammary bud. It is assumed this process happens in a highly coordinated process involving induction from the underlying ECM of the mesoderm, spatial patterning, cell proliferation, and differentiation. While formation of the milk line and primary bud are not observed in this model, remarkably, the highly coordinated events of induction, spatial patterning, cell proliferation, and differentiation are indeed captured. Moreover, this study represents the first live imaging to visualize this process in real time.

In vivo, the primary bud subsequently begins growth downward and branches, yielding secondary buds, which eventually develop into the mammary TDLU of the adult breast. Since the mesodermal cues this 3D hydrogel model are lacking, formation of the milk line and primary bud formation are bypassed. Instead, we observe the emergence of mesenchymal-like EMT’d cell emerging from growing organoids that support the development of TDLUs. In addition, we observe a unique feature of the non-classical EMT driving the bending of the collagen gel into a three-dimensional sphere with TDLUs that closely mirror those in the adult breast on the inside and surface cells on the outside. Furthermore, like the in vivo setting where alveolar buds invade into the matrix, we can visualize this invasion in real time. Therefore, despite bypassing the formation of a primary bud which creates the lactiferous ducts and the nipple, this 3D hydrogel model is still highly relevant for the study of human breast tissue development.

While the initial stages of organoid formation mirror early embryonic breast development, the subsequent stages diverge, offering an alternative, yet effective, path to achieve ductal branching and differentiation. This suggests that there are multiple valid ways to achieve the complex architecture of breast tissue. In addition, the ability of organoids to potentially bypass certain in vivo steps might offer advantages for studying breast development and for modelling disease in a controlled environment. This also underscores the importance of studying and understanding organoids as they can reveal alternative developmental processes and mechanisms not observed in vivo. For example the spiral cell movements in breast organoids, which have also been reported in branched lung organoids ^34^ remain to be studied and well understood. Thus, while this organoid model appears to mirror some of the events of ectoderm to a TDLU formation in its current form, it does lay the foundational structure upon which subsequent development stages, including ductal branching and lobuloalveolar development during puberty and pregnancy, could be incorporated and studied.

Second, this methodology facilitates the real-time study of cellular reprogramming and lineage plasticity opening new avenues for regenerative medicine and cancer research. Stem cell lineage reprogramming can be achieved simply by changing the elasticity and components of the matrix and after several weeks, the cells commit to their new lineage ^35^. Adult human breast tissue contains variant epithelial cells that can epigenetically reprogram from lineage-committed precursors in a process resembling iPSCs and demonstrate significant lineage plasticity, having the potential to differentiate into functional derivatives of all three germ layers (ectodermal, endodermal, and mesodermal) and even make skin ^36–38^. Here, we have observed the emergence of lineage divergent cells from growing organoids, that remain separate from the organoid parenchyma, function to bend and remodel the collagen hydrogel, and express markers of EMT and mesenchymal cells. We have also observed a prominent population of mesenchymal cells by scRNA-seq and functionally playing an active role in tissue architecture, assisting in the organoid structure’s integrity and stromal remodeling. Further verification and examination of mesenchymal cell origins in the hydrogel organoid model is warranted. While we have previously observed, using static microscopy imaging, that some organoids are surrounded by mesenchymal-looking cells (Supplemental Fig. S9c), their hyper-motility and participation in organoid maturation were only revealed through live imaging. The mesenchymal cells, with their pivotal role in the organoid’s structural integrity and stromal remodeling, warrant further exploration in subsequent studies.

Taken together, these observations made possible with this fine-tuned methodology offers a profound insight into the orchestrated events of human breast organoid formation, mirroring embryonic development stages. Single cells divide to establish a niche and progeny where one group of cells influences the development of another group of cells through close-range interactions. Spatial and temporal regulation, alongside cellular dynamics, contribute to the sophisticated architecture of organoids. This nuanced understanding can provide foundational insights into organ development, cellular and molecular mechanisms underlying tissue regeneration, and disease mechanisms.

### Limitations

While the presented technology offers a significant advancement, several limitations warrant consideration. Foremost, the model currently lacks the integration of certain stromal cells, most notably immune and vascular cells, which play crucial roles in tissue development, remodeling, homeostasis, and hormonal responses. Additionally, a prominent component of the breast is adipose tissue, which is absent in this model. This is a drawback, given the critical role of the adipose tissue in breast tissue endocrine functions. The model’s throughput, capped at 96 wells, limits the scale at which parallel experiments can be conducted, potentially hindering rapid screening and comparative studies. Moreover, while patient-derived cells enhance physiological relevance, they might introduce variability between experiments due to individual differences, posing challenges in standardization. Finally, as with any in vitro model, translating findings to in vivo conditions and clinical scenarios must be done with consideration, given the systemic factors and complex microenvironments that exist in a living organism.

## Methods

### Ethics statement

Primary tissues that would otherwise have been discarded as medical waste following surgery were obtained in compliance with all relevant laws, using protocols approved by the institutional review board at Maine Medical Center and Tufts Medical Center. All tissues were anonymized before transfer and could not be traced to specific patients; for this reason, this research was provided exemption status by the Committee on the Use of Humans as Experimental Subjects at the Massachusetts Institute of Technology, and at Tufts University Health Sciences (IRB# 13521). All patients enrolled in this study signed an informed consent form to agree to participate in this study and for publication of the results.

### Tissue processing and 3D culture

Tissues were partially dissociated with 1.5mg/ml collagenase (10103578001, Millipore Sigma) and 18μg/ml hyaluronidase (H3506, Sigma Aldrich), yielding small epithelial fragments, washed, and cryopreserved in 1 mL aliquots. Tissue fragments were further dissociated to single cells using 0.25% trypsin (25200056, Thermo Fisher Scientific) and 5 U/ml dispase II (4942078001, Roche). Hydrogels were seeded at a concentration of 500 cells per 200 μL gel (2.5 cells/μL). For hydrogel fabrication and seeding, cells were resuspended in 1.7 mg/ml collagen I (08-115, Millipore Sigma), 10 μg/ml hyaluronic acid (385908, Millipore Sigma) 40 μg/ml laminin (23017-015, Gibco), and 20 μg/ml fibronectin (F1056, Sigma Aldrich), pH 7.3. Gels were deposited in 4-well chamber slides (354104, Falcon) and incubated 1 hour at 37 °C for polymerization before adding MEGM medium ^10^. Hydrogels were cultured floating in serum-free mammary epithelial growth media (MEGM) in 4-well chamber slides. MEGM consisted of basal medium (M171500, Thermo Fisher Scientific) supplemented with 52μg/ml bovine pituitary extract (13028014, Thermo Fisher Scientific), 10ng/ml hEGF (E9644, Sigma Aldrich), 5μg/ml insulin (I9278, Sigma Aldrich), 500ng/ml hydrocortisone (H0888, Sigma Aldrich), 1% v/v GlutaMAX (35050061, Thermo Fisher Scientific), and 1% v/v penicillin-streptomycin antibiotics (15140122, Thermo Fisher Scientific).

### Immunofluorescent Staining of Hydrogels

Hydrogels were fixed with 4% paraformaldehyde (15710-S, Electron Microscopy Sciences) at room temperature for 30 minutes and washed in PBST, comprised of 1X PBS (46-013-CM, Corning) and 0.05% Tween 20 (P7949, Millipore Sigma). Samples were then permeabilized in 0.1% TritonX-100 (X100, Sigma Aldrich) in PBST and incubated with blocking buffer comprised of PBST with 3% BSA (BSA-50, Rockland Immunochemicals) and 10% goat serum (G9023, Millipore Sigma) for 2 hours at room temperature. Hydrogels were then stained with the appropriate primary antibody in blocking buffer at 4°C overnight. Samples were washed with PBST and incubated with secondary antibody for 2 hours at room temperature, and DAPI for 30 minutes at room temperature. Primary antibodies used in this study were: E-cadherin (ab1416, Abcam, Clone HECD-1, 1:100), CK14 (RB-9020, Thermo Fisher, 1:300). Secondary antibodies used in this study were: goat-anti-mouse-AF555 (A21424, Life Technologies, 1:1000), goat-anti-rabbit-AF488 (A11008, Life Technologies, 1:1000). Alexa fluor 546-conjugated anti rabbit (A11010, Invitrogen, 1:500) and Alexa fluor 546-conjugated anti sheep (A21098, Invitrogen, 1:500). Dyes and probes used for immunofluorescence: Phalloidin-AF647 (A22287, Life Technologies, 1:250 of 400x stock), DAPI (D1306, Life Technologies, 5μg/ml), Hoechst 33342 (H3570, Thermo Fisher, 1:1000).

### FFPE Patient Tissue Staining

Patient tissue was fixed in 10% buffered formalin and paraffin-embedded in tissue blocks which were serially sectioned. Slides were baked at 55°C for 45 min, then rehydrated by sequential incubation in Xylene, 100% ethanol, 95% ethanol, 70% ethanol, and 50% ethanol for 3 minutes each, then 10 minutes in PBS. Antigen retrieval was done in sodium citrate buffer in a steamer for 20 minutes. Slides were washed in TBS-T comprised of 1X TBS (Trizma base, T4661, Millipore Sigma; NaCl, 71376, Millipore Sigma) and 0.25% TritonX-100 and then in blocking buffer, comprised of TBS-T with 10% goat serum and 1% BSA, for 2 hours at room temperature. Slides were stained with appropriate primary antibody in blocking buffer at 4°C overnight. Samples were washed with TBS-T and incubated with secondary antibody for 2 hours at room temperature, and DAPI for 30 minutes at room temperature. Slides were washed in tap water and covered with mounting medium and glass coverslips. Primary antibodies used in this study were: E-cadherin (610182, BD Biosciences, Clone 36, 1:50), CK14 (RB-9020, Thermo Fisher, 1:50), Secondary antibodies used in this study were: goat-anti-mouse-AF555 (A21424, Life Technologies, 1:1000), goat-anti-rabbit-AF488 (A11008, Life Technologies, 1:1000). Dyes and probes used for immunofluorescence: DAPI (D1306, Life Technologies, 5μg/ml).

### Live imaging

Primary single cells, isolated as described above, were incubated with the cell tracking dye Cytopainter Green (ab138891, Abcam, 1:1000) for 30 minutes and then were washed and seeded at a concentration of 100 cells per 20 μL hydrogel. For gel fabrication, 20 μL hydrogel drops were deposited onto the center wells of a 96-well plate (3603, Corning). Gels were allowed to incubate for one hour at 37°C until fully polymerized. 80μL of MEGM was then added to each well and gels were gently lifted off the well surface with a pipette tip. Cultures were immediately placed in a pre-warmed incubator chamber (Okolab Inc) enclosed over a Nikon Eclipse Ti2-AX confocal microscope (Nikon). Images of selected points were collected starting immediately after the addition of media, and every 30-45 minutes after, in both brightfield and with A488 laser at 4x magnification and 2.5x zoom across nine z-positions. 20-40 μL media was added to the culture twice a week. Cultures were live imaged for 18-21 days. Analysis and production of videos across locations and timepoints was performed using NIS-Elements (Nikon) and Premiere Pro (Adobe) software.

### Transduction of primary cells in hydrogels

Lentiviral particles were produced in 293T cells (seeded at 6*10^6^ cells per plate), grown in 10 cm plates with 6 mL of Optimem (Gibco, 31985070). 293T cells were transfected using Fugene 6 (20 μL, Promega, PAE2691), with the following plasmids: 2 µg pCMV-VSV-G, 4 µg pCMV-dR8.2-dvpr and 4 μg of expression plasmid. Media was collected from plates on day 2 and 3 after transfection, filtered through a 0.45 μm filter and used immediately or stored at -80°C in 1 mL aliquots. Primary mammary epithelial cells were placed in an ultra-low attachement 24 well plate (Corning, #3473) at a concentration of 10^5^ cells per well, in 150 μL MEGM supplemented with protamine sulfate (5 μg/mL). Lentiviral vectors containing LeGO-V2 (Addgene plasmid #27340 ^19^) and pBOB-EF1-FastFUCCI-Puro (Addgene plasmid # 86849 ^22^) were added in volumes of 50 μL to bring to total volume in each well to 200 μL. Cells were centrifuged at a speed of 1000 x g and at a temperature of 30°C for 2 hours and then incubated overnight. Cells were washed, passed through a 40 μM cell strainer (VWR, 76327-098), and counted prior to seeding in the hydrogel.

### Cell Tracking and Data Acquisition

#### Individual cell tracking

Using FIJI software ^39^, 15 individual cells across eight videos of live confocal imaging were tracked, depicting the development of organoids from single cells. The x/y coordinates of each cell were manually annotated frame-by-frame to generate precise tracking data. The speed of each cell was calculated for each segment by measuring the displacement over the time interval between successive frames. The frame rate of the video recordings was 20-35 minutes (a precise rate was known for each frame) allowing a high temporal resolution. Visualization of cell tracks and speed measurements was carried out using R (Version 4.3.3) employing the dplyr, ggplot2, and readxl packages. Adjustments were made to the cell coordinates, anchoring the starting point at (0, 0) to facilitate a clearer visualization of movement paths. The positioning of each cell was determined in each frame, and speed was deduced from the elapsed time between frames. The correlation coefficient (CC) between the X and Y coordinate values was calculated to quantify the conformance of the track to a linear path, where a CC closer to 1 or -1 indicated a greater degree of conformance to a linear path. Low or Near Zero CC suggests no clear linear trajectory, indicative of random or highly erratic movement without a preferred direction.

#### Cell population tracking

Cell populations were tracked in two live imaging videos. FIJI software and the FIJI plugin TrackMate ^39,40^ were used to automatically identify cells across frames with LoG-based (Laplacian of Gaussian) detectors. To ensure consistent tracking, frame shifts in the video were corrected. The x/y coordinates of detected events, representing the spatial positions of cells in each frame of the video, were analyzed using R software ^41^. A focal point was defined for each video, and the distance of each cell from the focal point was calculated as 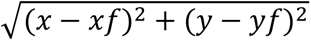, where *xf* and *yf* are the coordinates of the focal point. Cells were categorized into distance ranges based on their calculated distances. Data visualization was performed using ‘ggplot2’ in R.

#### Tracking organoid growth

Videos from live imaging experiments, initiated on days 1, 8, and 9, were processed at a rate of five frames per second and extracted as individual frames. Each set of frames was analyzed using “ilastik”, a neural network designed for prediction of tissue versus background ^42^. A training set of 8 frames per video was employed, allowing ilastik to generate probability maps indicating tissue presence in each frame. These maps were subsequently processed using CellProfiler ^43,44^, where images were converted to grayscale, background noise was reduced, and any remaining debris was suppressed while enhancing the edges of the organoids. CellProfiler was also utilized to identify primary objects (organoids) and track their development by measuring the area they occupied in each frame. To maintain accurate measurements across time, the number of frames per 6-hour interval was calculated by dividing the total number of frames by the duration of organoid development. Measurements for each 6-hour interval were averaged together. To account for variability in pixel counts from CellProfiler output, the area measurements for each organoid were normalized to the first accurate measurement of each organoid’s area, yielding fold change in area during development. These normalized values were scaled to provide absolute measurements in microns. Lines of best fit were determined by plotting both Gompertz function for sigmoidal growth and log transformed linear regression for exponential growth in PRISM 10. Parameters for the Gompertz model were configured to allow up to 1000 iterations to ensure convergence, with no special handling of outliers applied to maintain data integrity. The YM value (asymptotic maximum) was unconstrained, allowing the model to estimate the maximum response based on the data. The initial value, Y0, was constrained to be greater than 0, and the growth rate constant, K, was constrained to be greater than 0 to ensure a positive growth rate. A 95% confidence interval was applied to all estimated parameters, providing an estimate of the precision and reliability of the model’s predictions. Pearson correlations were calculated to examine the relationship between organoid size and growth rate with an additional two-tailed t-test to assess the statistical significance of the linear regression models. The goodness of fit for each regression model was assessed using the R-squared (R²) value. An R² value of “NA” was recorded in instances where the linear regression model could not be calculated within the specified parameters, indicating that the model failed to converge or that the data did not fit the Gompertz function adequately.

### Single cell RNA sequencing & analysis

#### Sample preparation, sequencing, and analysis

Five patient samples were plated into 12 hydrogels each, at 4,000 cells/gel. Samples grew for 19 days, and then were dissociated into single cells and pooled. Cells were dissociated from the hydrogel by 30 minutes incubation with collagenase (500 μg/mL) in PBS, followed by incubation in trypsin for 5 minutes. Cells were washed and resuspended in PBS, strained through a 40μm cell strainer, and counted using a cell counter. Sample preparation and cDNA library generation was done using 10X Chromium Next GEM Single Cell 3’ kit v3.1 following manufacturer’s protocol. Libraries were sequenced on the Illumina NovaseqX platform, Novogene.

Raw paired end reads were be demultiplexed and aligned to the GRCh38 human genome using Cell Ranger 8.0.0. Once reads are aligned, the data was analyzed using Seurat v5^29^ for data normalization and feature selection. Filtering eliminated genes detected in <3 cells, and removed cells with <200 or > 7500 genes and with mitochondrial content >15%. Data was normalized, and variable features were identified using the FindVariableFeatures function. Data was then scaled and PCA analysis ws performed. Clustering was done using the FindClusters function, using a resolution of 0.5 and the first 10 dimensions of the principle component analysis. Clustered cells were visualized by UMAP embedding using the default settings in Seurat. FindAllMarkers function was used to identify the top differential genes for each cluster. Pseudotime analysis was performed using Slingshot v2.12.0. For pseudotime analysis, each branch represents a path for one estimated lineage which are based on transcriptomic similarities. The green dot represents the pseudotime 0 start point of the analysis. Clusters were annotated based on the differential gene expression of canonical markers, as well as pseudotime analysis.

#### Integrated Analysis

Previously published scRNA-seq data from traditional organoids grown in Matrigel^45^, healthy primary breast tissue^12^, as well as the scRNA-seq data generated from this study were analyzed. Raw paired end reads were be demultiplexed and aligned to the GRCh38 human genome using Cell Ranger 8.0.0. Once reads are aligned, the data was analyzed using Seurat v5^29^ for data integration, normalization, and feature selection. Briefly, 10X files were loaded and Seurat objects were generated for each data set. Filtering removed cells with < 200 or >7500 genes and with mitochondrial content >7.5%. Genes detected in less than 3 cells were dropped from analysis. The Seurat objects were merged using the merge function, and the data was then normalized by multiplying transcripts by a factor of 10,000 and then log-transforming the data. Variable features used for analysis were identified by using the FindVariableFeatures function at default settings. Data was then scaled and PCA analysis was run. Integration was done with the CCAIntegration method using the IntegrateLayers function in Seurat. Cells were then clustered using K-nearest neighbor (KNN) graphs and the Louvain algorithm using the first 15 dimensions from the dimensional reduction analysis. Clusters were called using the FindClusters function with a resolution of 0.3, and 17 distinct cell clusters were identified. Clustered cells were visualized by tSNE embedding using the default settings in Seurat. The layers of the integrsted Seurat object were then joined using the JoinLayers function. To identify differentially expressed genes between cell clusters, we utilized the FindAllMarkers function to identify features detected in >10% of a cell cluster compared to all other cells.. Pathway analysis to identify enriched biological pathways associated with differentially expressed genes was done using established databases, such as PanglaoDB ^30^. The top differentially expressed markers were used to determine gene expression location.

### Hormone treatment

Hydrogels were seeded with single primary cells from patient samples: pre-menopauseal (n=5 patient sampels, 4 hydrogels per patient), post-menopausal (n=5 patient sampels, 4 hydrogels per patient) and patients undergoing female-to-male transition (F-M, n=5, 4 hydrogels per patient), at a concentration of 1000 cells per hydrogel. Hydrogels were deposited and cultured floating in 4-well chamber slides, as described above (Tissue Processing and 3D Culture). Hormone supplements were dissolved in 0.5% DMSO. For hormone treatment experiments, media was supplemented with either 0.5% DMSO (317275, Millipore Sigma), or 1nM 17β-estradiol (E2758, Sigma) and 1 nM progesterone (P0130, Sigma). Hydrogels were cultured in hormone treatment media for 19 days with media changes occurring every 2-3 days. Organoid number and morphology were analyzed on day 19.

### Flow Cytometry

Single cells dissociated from patient tissues as described above were washed with PBS with 2% FBS and stained with antibodies at 4°C for 60 minutes. Cells were then washed and resuspended in PBS with 2% FBS for flow cytometry. All data acquisition was done using an LSR II (BD Biosciences) and analyzed using FlowJo software (TreeStar). Antibodies used for flow cytometry: CD49f-FITC (555735, BD Biosciences, 1:20), EpCAM-APC (347200, BD

Biosciences, 1:20).

## Data and code availability

- This paper analyzes two existing scRNA-seq datasets, publicly available through the following GEO accession numbers: GSE180878 ^46^ ; GSE162296 ^15^.
- Original scRNA-seq data for this paper was deposited to GEO, and is available through the following accession number: GSE274346.
- Any additional information required to reanalyze the data reported in this paper is available from the lead contact upon request.

## Acknowledgments

We would like to extend our sincere gratitude to Jean-Yves Tinevez from the Institut Pasteur, Université Paris Cité, for his invaluable advice and support. His expertise and guidance on the cell tracking FIJI plugin TrackMate ^39,40^, was instrumental to the success of this work.

This research was supported by the following: NIGMS (7R01GM124491, CK), DOD-BCRP (W81XWH2010018, GR), Breast Cancer Research Foundation (CK) and FTC Breast Cancer Foundation (CK).

## Author Contributions

GR, NCT, CJT, MEP and CK conceived the project and designed experiments. GR, NCT, CJT, MEP and KM performed experiments. GR, NCT, CJT, MEP and CK analyzed results GR, NCT, CJT, MEP and CK wrote the manuscript with contributions, editorial review, and approval from all authors.

## Supplemental Video Legends

**Supplemental Video 1:**

This video demonstrates the full process of organoid development, showcasing the four phases: induction, patterning, morphogenesis, and maturation. This video demonstrates Type II induction, where cells start from a close-packed spheroid. Individual elongated, spindle-shaped cells are ejected from the spheroid colony, initiating the patterning phase. The branch subsequently grows in the direction of the preceding cell. The final maturation phase displays the development of terminal ductal-lobular units (TDLUs). Following the Maturation phase, the gel contracts, and the mature organoid is surrounded by a mesemchyme.

**Supplemental Video 2:**

This video captures the mesenchymal-like cells’ dynamic interaction with the collagen matrix during organoid development. These cells actively contract and manipulate the collagen matrix, leading to the condensation of the hydrogel. This video also illustrates the synchronized, wave-like sinusoidal pattern of mesenchymal cell movement.

**Supplemental Video 3:**

Live imaging demonstrating Type I induction, showing cells initially exhibiting minimal movement, then progressing to entrer the patterning phase, with individual cells migrating and forming the initial blueprint of the organoid.

**Supplemental Video 4:**

This video demonstrates Type I (organoid on top) and Type II (organoid below) inductions, and highlights the patterning phase in detail. During patterning, a fluorescently labeled cell migrates along the axis of the extending branch, returning to the main body, and then moving back to the invasion front. This dynamic movement lays the branch pattern for the subsequent morphogenesis phase. Spiral cell rotation is observed in the terminal acini. Scale bar = 100 μm.

**Supplemental Video 5:**

Live imaging of organoid development reveals the expansion of mesenchymal-like cells from the central organoid point. Arrows indicate organoid regions from which mesenchymal-like cells appear to exude. The video captures the proliferation and scattering of these cells, actively remodeling the collagen matrix. Following the Maturation phase, the gel contracts, and the mature organoid is surrounded by a mesemchyme.

**Supplemental Video 6:**

Live imaging video of organoid formation from single cells to mature TDLUs. The video provides a view of the different stages, including induction, patterning, morphogenesis, and maturation, highlighting the coordinated cellular movements and interactions throughout the process. Following the Maturation phase, the gel contracts, and the mature organoid is surrounded by a mesemchyme.

**Supplemental Video 7:**

Cells transduced with the FastFUCCI vector are tracked during organoid development, displaying changes in the cell cycle. Cells fluorescing red are in the G1 phase, while those fluorescing green are actively dividing in the S, G2, or M phases. The video illustrates the dynamic repositioning and movement of cells within the growing TDLU during different phases of the cell cycle, and spiral movement of the cells in the terminal acini. Scale bar = 100 μm.

**Supplemental Video 8:**

Emergence and distribution of mesenchymal-like cells. The video highlights the proliferation and radial expansion from the central point, with cells actively participating in matrix remodeling and organoid shaping. Scale bar = 100 μm.

**Supplemental Video 9**:

High-resolution imaging of organoid maturation, illustrating the coordinated movements and rearrangements of cells. The video captures the spiral rotation in terminal acini and longitudinal migration along the ducts. Scale bar = 100 μm.

**Supplemental Video 10:**

Live imaging of organoid development. Arrows indicate organoid regions from which mesenchymal-like cells appear to exude.

**Supplemental Videos 11-19:**

Time-lapse videos of organoid development by breast primary cells derived from patient samples were, captured at 30-minute intervals, monitoring organoid morphological changes and growth dynamics during the end of Patterning through Morphogenesis and Maturation phases.

---

**Supplemental Fig. S1:**
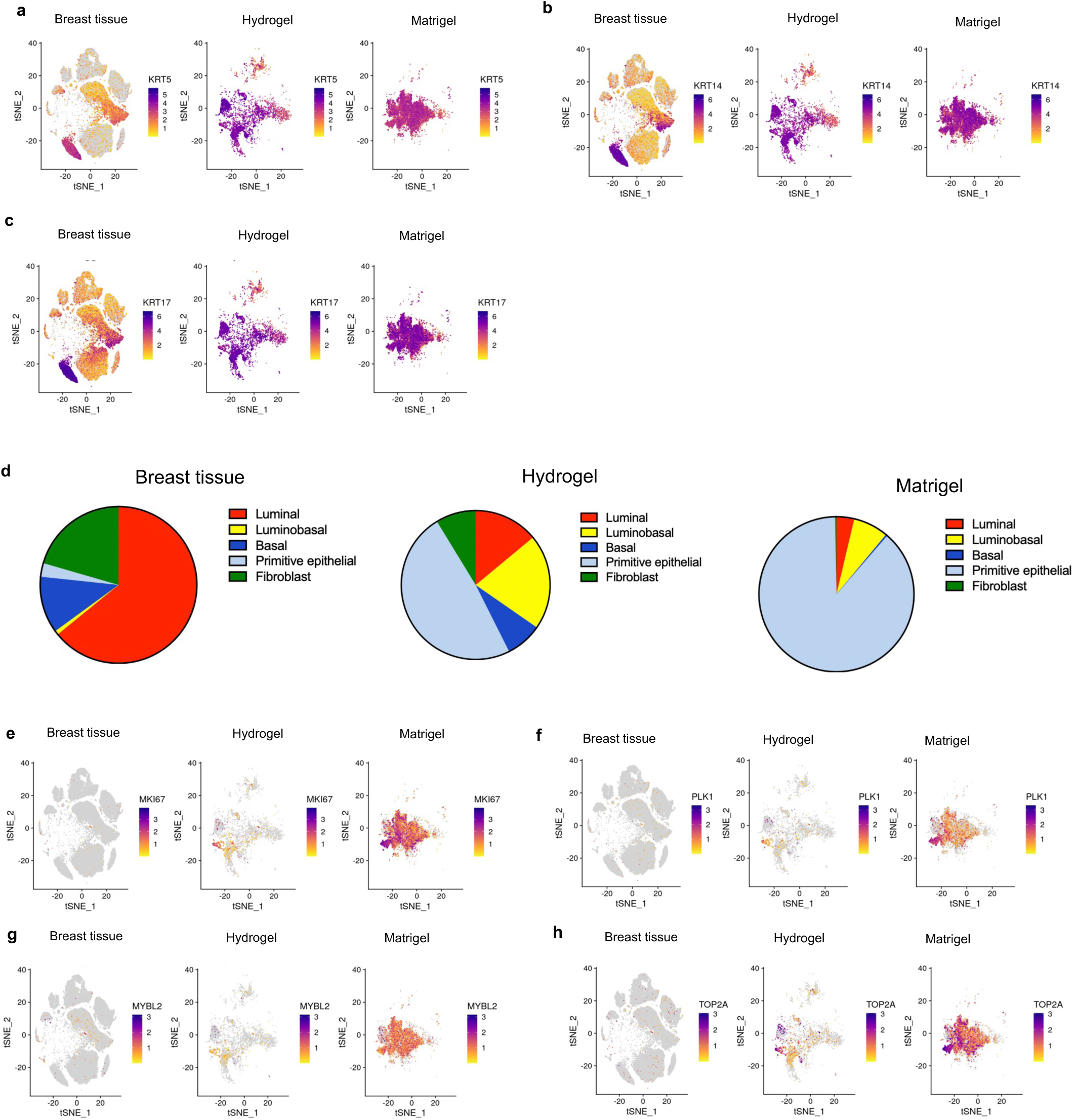
scRNA-seq analysis of breast tissue and organoids grown in ECM hydrogel or in Matrigel. **a-c**, Expression patterns highlighting basal markers KRT5, KRT14, KRT17. **d,** Proportion of the 5 major cell types shared across human breast tissue, hydrogel organoids, and materigel organoids. Percentages were normalized after the removal of cell populations not shared across the three conditions. **e-h,** Expression patterns highlighting basal markers MKI67, PLK1, MYBL2 and TOP2A

**Supplemental Fig. S2:**
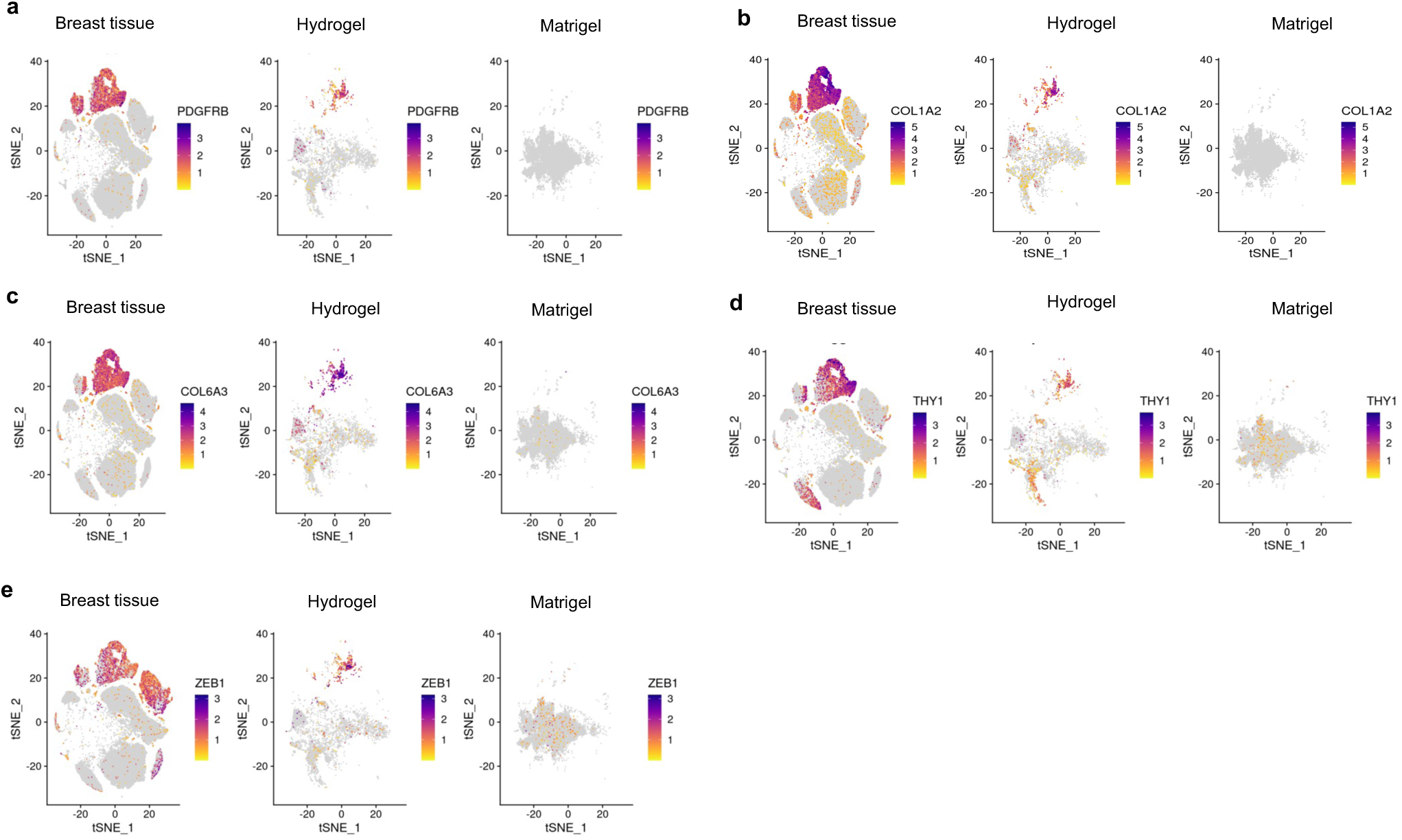
scRNA-seq analysis of breast tissue and organoids grown in ECM hydrogel or in Matrigel. **a-e**, Expression patterns highlighting stromal fibroblast markers PDGFRB1, COL1A2, COL6A3, THY1, and ZEB1.

**Supplemental Fig. S3:**
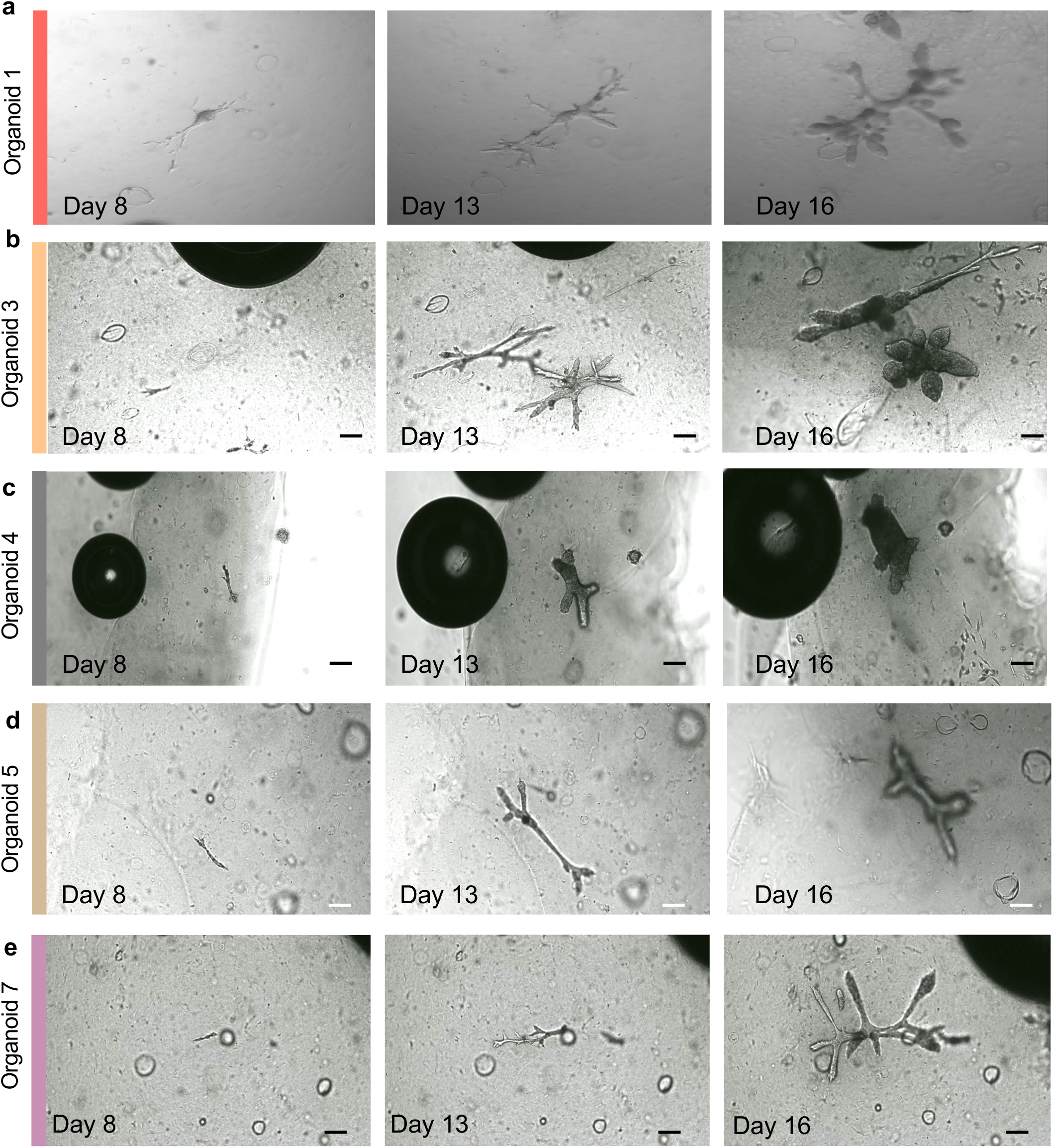
Organoid growth. (**a-e):** Representative images of organoids from the time-course plot, captured on days 8, 13, and 16, highlighting the morphological changes and growth progression of individual organoids over time

**Supplemental Fig. S4:**
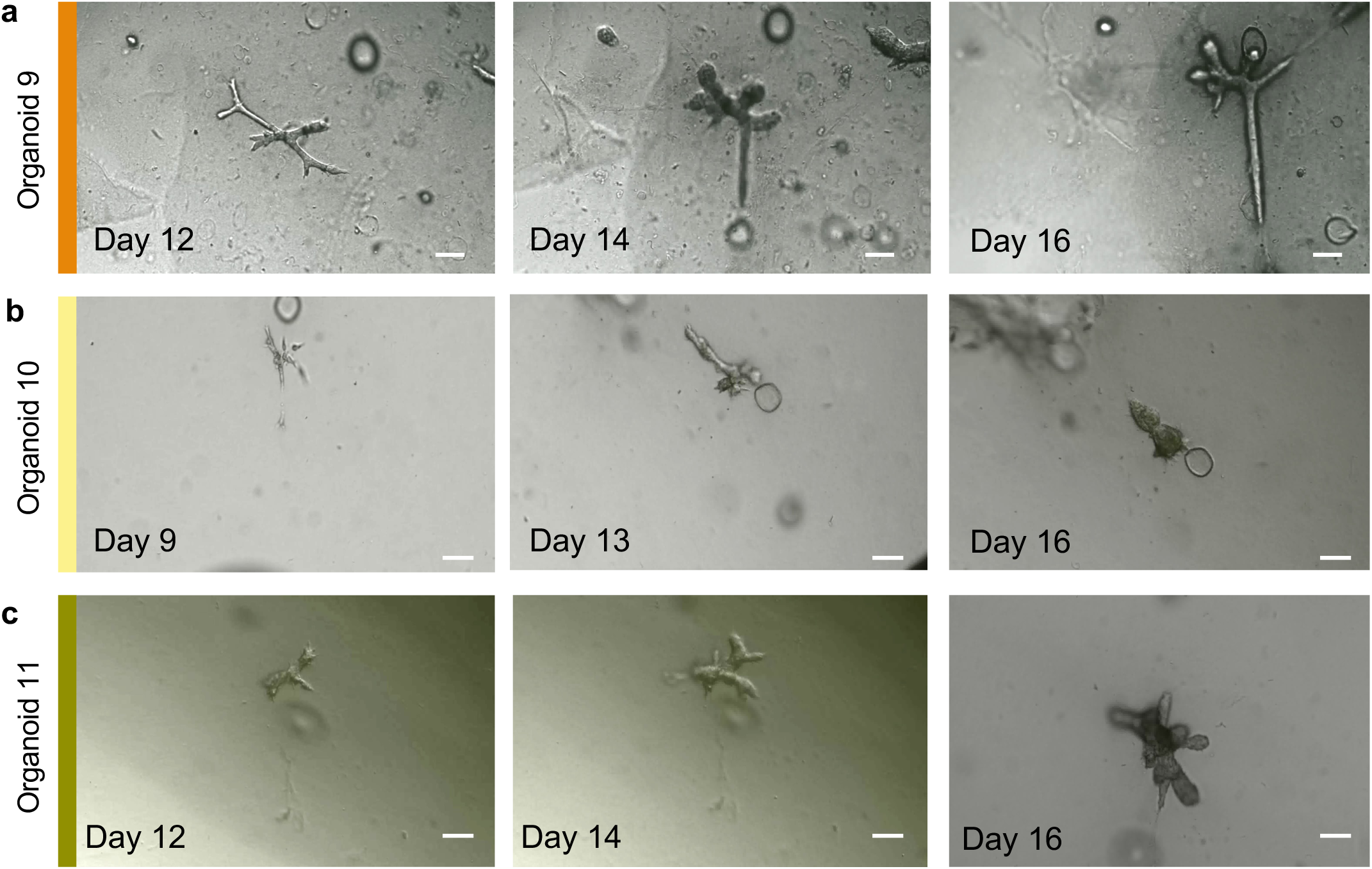
Organoid growth. (**a-c):** Representative images of organoids from the time-course plot, captured on days 8, 13, and 16, highlighting the morphological changes and growth progression of individual organoids over time

**Supplemental Fig. S5:**
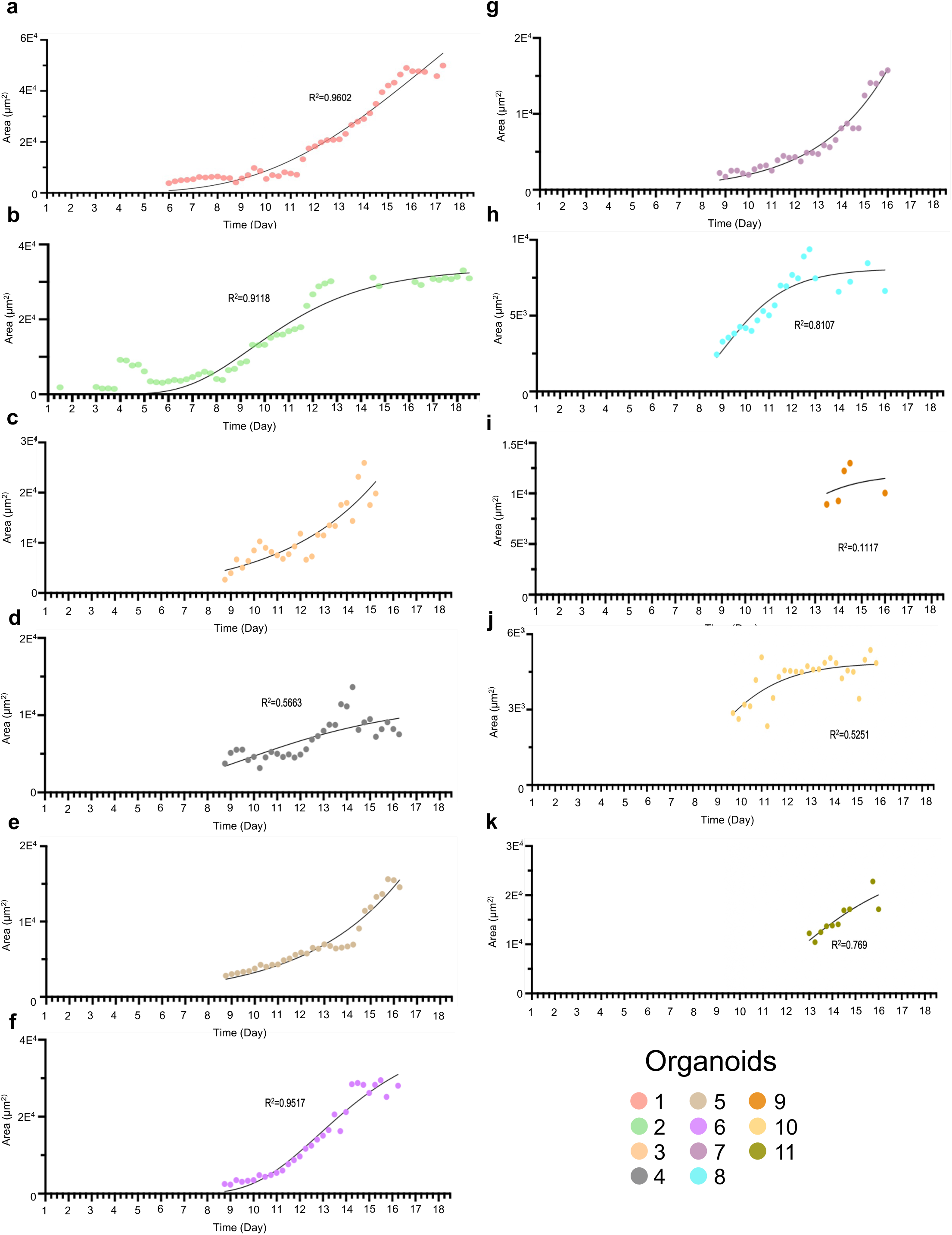
Linear regression analysis of individual organoid growth using the Gompertz function. **(a-k):** Sigmoidal growth patterns of the organoids, characteristic of some biological growth processes. Each panel represents the growth curve of a single organoid, highlighting the variability in growth dynamics among the different organoids.

**Supplemental Fig. S6:**
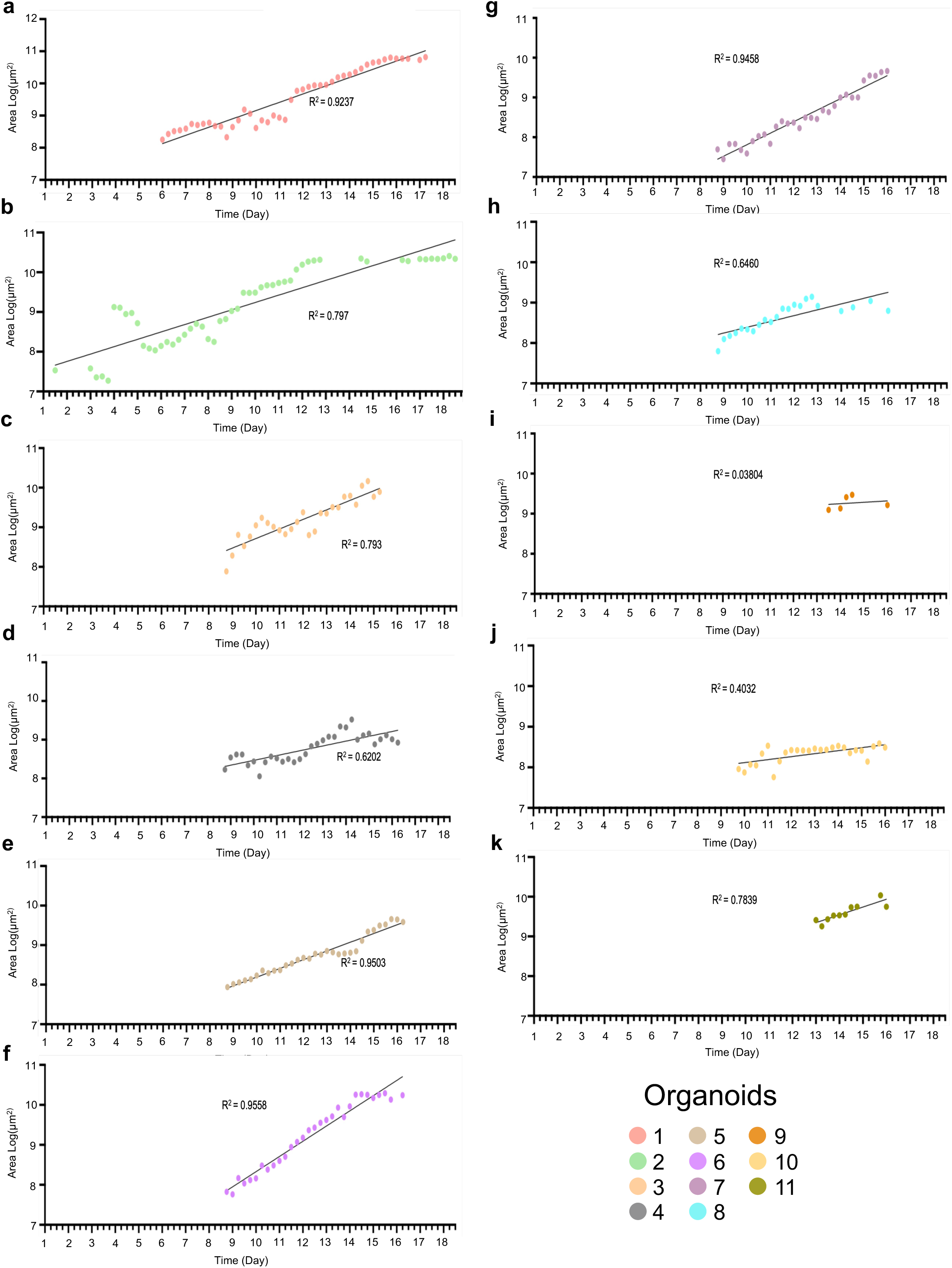
Linear regression analysis of individual organoid growth (using log-transformed data and linear regression. **(a-k):** Exponential growth patterns of the organoids analyzed by transforming the growth data to a logarithmic scale and fitting a linear regression model. Each panel shows the growth trajectory of a single organoid, demonstrating how the exponential growth model fits the observed data for each organoid.

**Supplemental Fig. S7:**
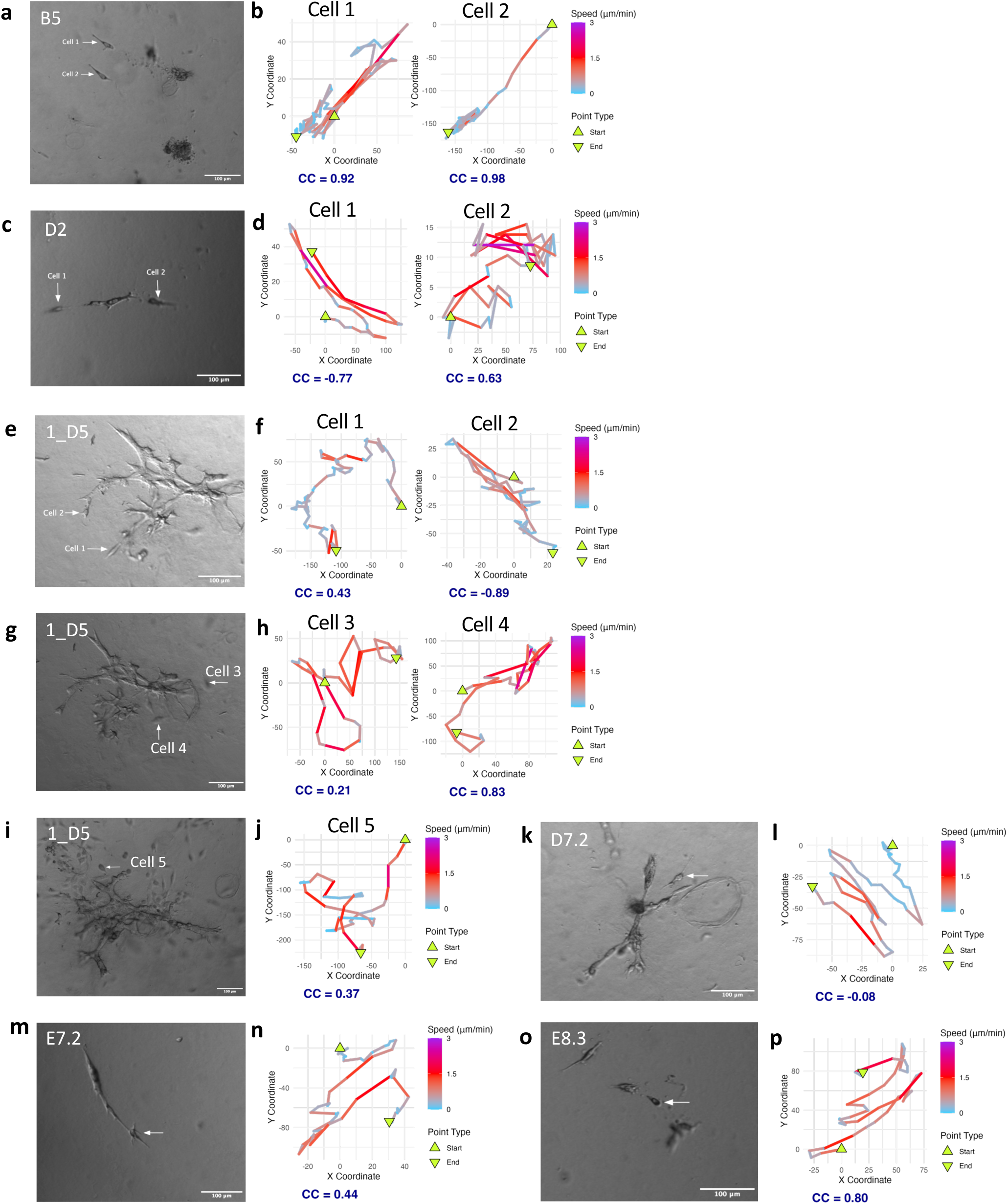
Analysis of Cell Movement Patterns and Path Conformance. **(a, c, e, g, i, k, m, o):** Single frame image from live imaging video, depicting tracked cells within the organoid culture. Scale bars = 100 µm. **(b, d, f, h, j, l, n, p)**: Corresponding trajectory plots. The color gradient indicates the speed of the cells. Triangles denote the starting and ending points of the tracks. The correlation coefficient (CC) between the X and Y coordinate values is denoted for each cell track.

**Supplemental Fig. S8:**
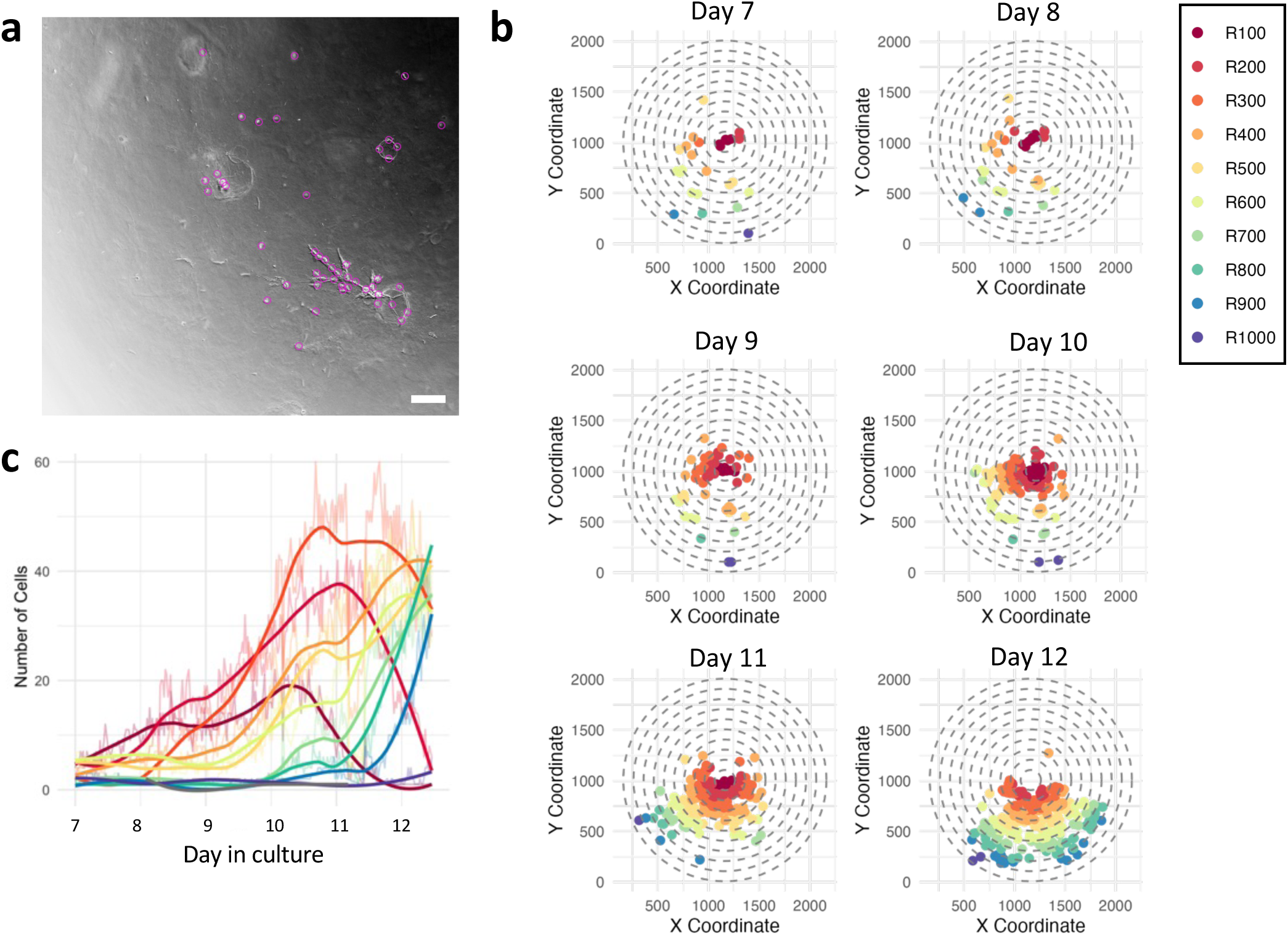
Tracking cell populations demonstrates radial expansion from organoid origin point. **a,** Representative frame from Supplemental Video 5 (inverted bright field). Purple circles mark spots detected by the cell tracking software (TrackMate). Scale bar = 100 μm. **b,** Plots showing the spatial distribution of cells at various distances from the center over time. Each plot represents a different time point. Cells are color coded corresponding to their distances from the center, as indicated in the legend on the right (e.g., R100 indicates distance of 100 μm from the center). **c,** Graph depicting the number of cells over time for each distance category from the center. Colored line corresponds to distance categories, matching the plots in ‘b’. Raw data lines are shown in the background and smoothed lines in the foreground.

**Supplemental Fig. S9:**
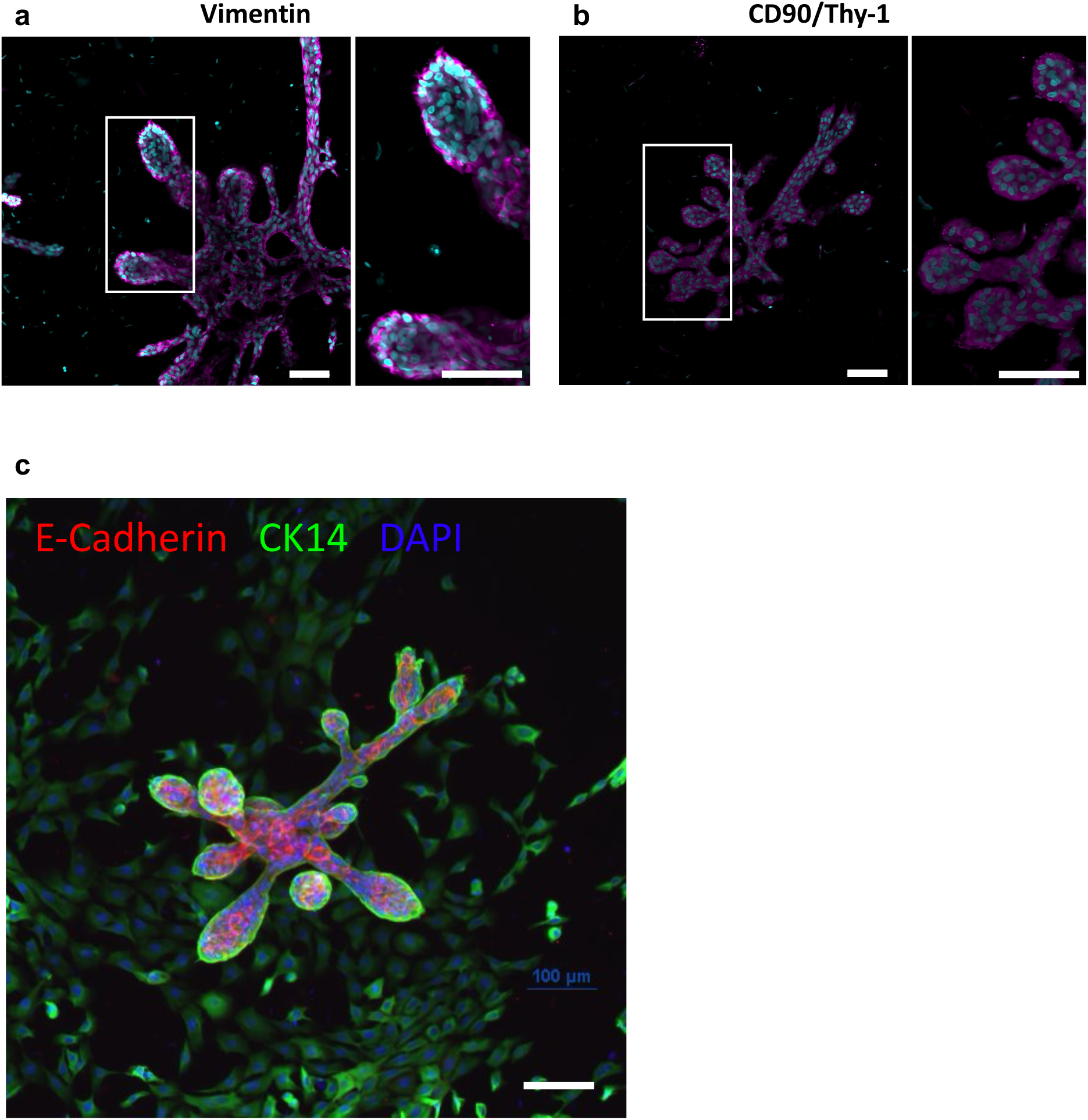
Immunofluorescence staining of primary breast organoid cultures demonstrating the expression of mesenchymal markers. **a,** Vimentin expression in basal regions of epithelial organoids. **b,** CD90/Thy-1 expression in basal regions of epithelial organoids. Magenta: protein markers; cyan: Hoechst nuclear staining. Right images in (a) and (b) represent higher magnification images of the regions indicated by the white boxes in left image. **c,** CK14 and E-cadherin expression in epithelial orgnaoids and surrpunding mesenchymal cells. Scale bars = 100 µm.

**Supplemental Fig. S10:**
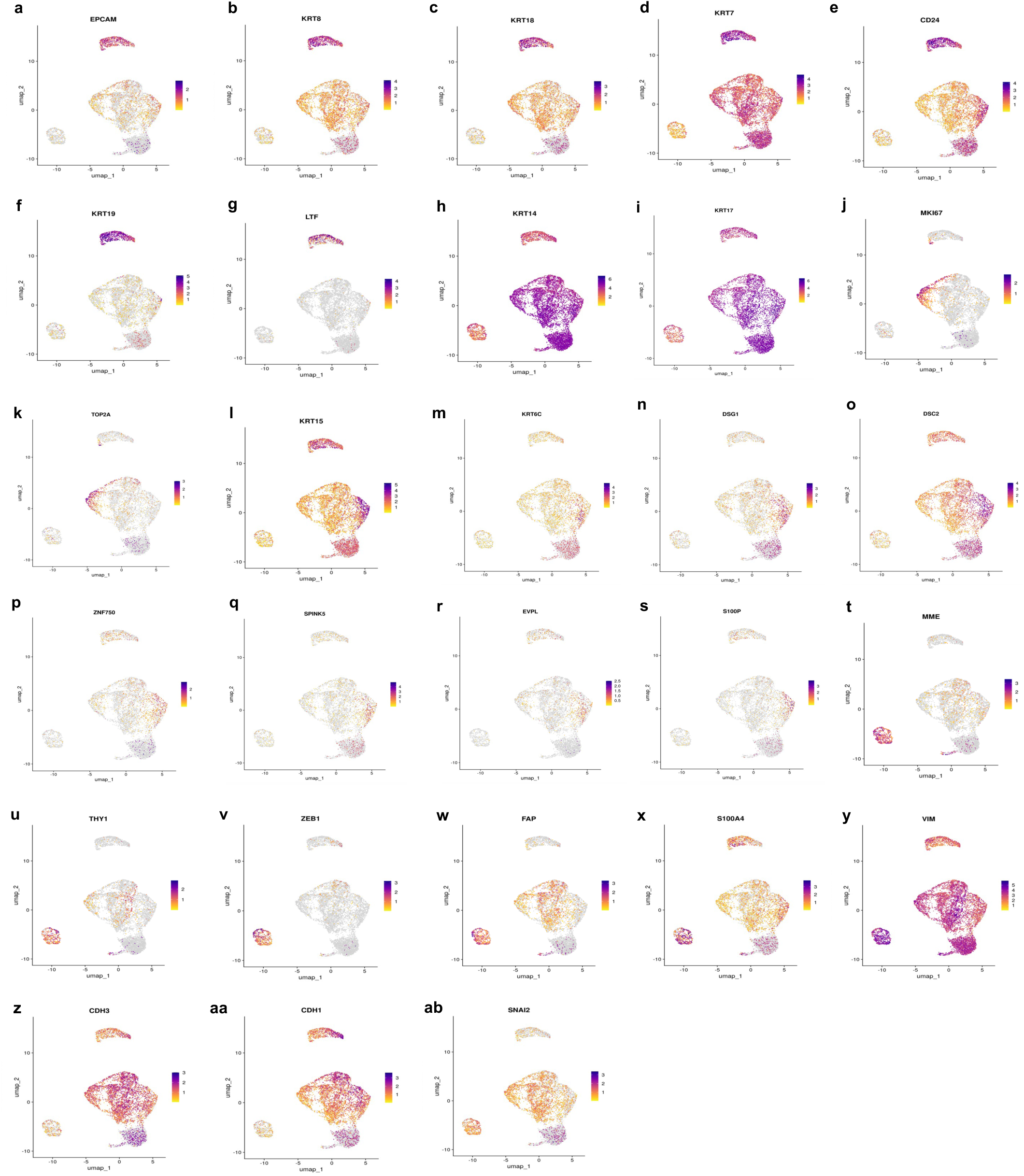
scRNA-seq analysis of organoids grown in ECM hydrogel. **a-g,** Expression pattern of luminal markers EPCAM, KRT8, KRT18, KRT7, CD24, KRT19 and LTF. **h-i,** Expression pattern of basal markers KRT14 and KRT17. **j-k,** Expression pattern of proliferative markers MKI67 and TOP2A. **l-s,** Expression pattern of genes related ot epidermal differentiation and keratinocyte function: KRT15, KRT6C, DSG1, DSC2, ZNF750, SPINK5, EVPL, and S100P. **t-x,** Expression pattern of mesenchumal markers MME, THY1, ZEB1, FAP, and S100A4. **y-ab**, Expression pattern displaying overlap between epithelial and mesenchymal markers: Vimentin, CDH3, CDH1, and Slug.

**Supplemental Fig. S11:**
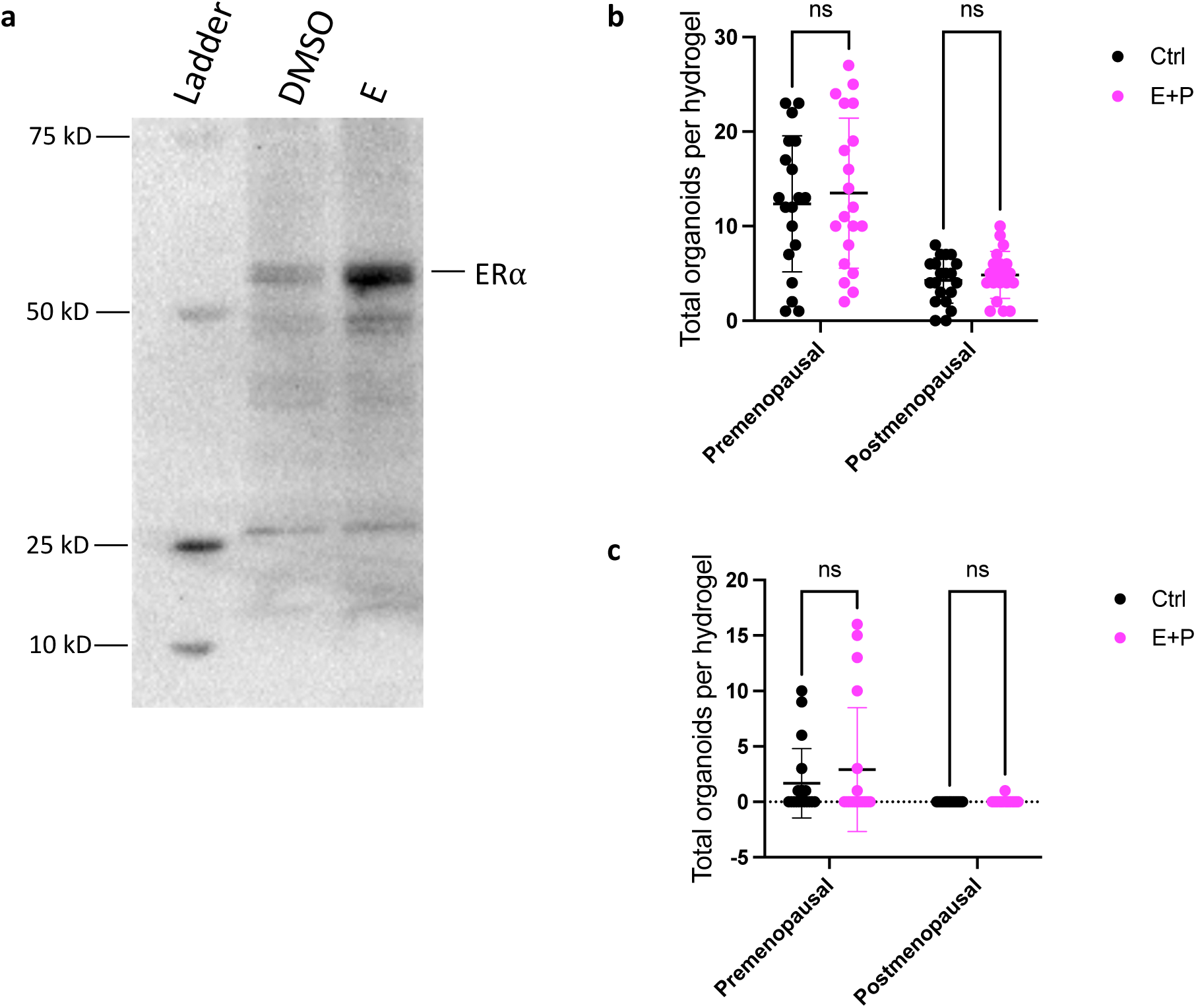
Effect of estrogen and progesterone treatement on organoid development and morphology. **a,** Western blot analysis of ERα after hydrogel growth with 1 ng/mL E2 for 21 days. **b,** Total organoids per hydrogel formed by premenopausal and postmenopausal patient samples treated with estrogen and progesterone (E+P), or vehicle control. **c,** Ductal-lobular organoids per hydrogel formed by premenopausal and postmenopausal patient samples treated with estrogen and progesterone (E+P), or vehicle control. (one way ANOVA with Sidak’s multiple comparisons test).

